# Modelling cortical laminar connectivity in the macaque brain

**DOI:** 10.1101/2020.07.19.210526

**Authors:** Ittai Shamir, Yaniv Assaf

## Abstract

In 1991, Felleman and Van Essen published their seminal study regarding hierarchical processing in the primate cerebral cortex. Their work encompassed a widescale analysis of connections reported through tracing between 35 regions in the macaque visual cortex, extending from cortical regions to the laminar level. In this work, we revisit laminar-level connectivity in the macaque brain using a whole-brain MRI-based approach. We use multi-modal ex-vivo MRI imaging of the macaque brain in both white and grey matter, which are then integrated via a simple model of laminar connectivity. This model uses a granularity-based approach to define a set of rules that expands cortical connections to the laminar level. Different fiber tracking routines are then examined in order to explore the ability of our model to infer laminar connectivity. The network of macaque cortical laminar connectivity resulting from the chosen routine is then validated in the visual cortex by comparison to findings from Felleman and Van Essen with an 83% accuracy level. By using a more comprehensive definition of the cortex that addresses its heterogenous laminar composition, we can explore a new avenue of structural connectivity on the laminar level.

## Introduction

In 1991, Felleman and Van Essen published their influential study regarding hierarchical processing in the primate cerebral cortex (Felleman and Van Essen 1991). The study encompassed a widescale analysis of cortical connections in the macaque brain, which have been reported using histological tracing. The connections link 35 cortical regions in the macaque visual cortex, extending from the regional level to the laminar level. Since then, their work has propelled the study of the interconnectedness of the visual cortex and has become a significant milestone in the field of structural connectomics.

Over the past two decades, great progress has been made in the field of MRI imaging of white matter connectivity (Van Essen et al. 2013, Setsompop et al. 2013). Since then, neuroimaging research of brain connectivity has focused on three principal types of connectivity: structural, functional and effective connectivity (Park and Friston 2013).

Currently, one of the major limitations of structural connectivity in general, and tractography particularly, lies in the lack of information regarding the laminar end points of white matter tracts inside the cortical grey matter (Jbabdi and Johansen-Berg 2011). In other words, the field of connectomics is currently limited by the definition of the cortex as a homogenous unit, focusing on transverse partitioning into regions and ignoring radial partitioning into laminar components. The integration of microstructural information regarding cortical properties and macrostructural connectomics could pose an exciting development in the field, by offering a multi-scale micro-level connectome (Johansen-Berg 2013).

In recent years, promising strides have also been made in the field of MRI imaging of the cortical laminar composition of the cortex (Clark et al. 1992, Barbier et al. 2002, Duyn et al. 2007, Barazany and Assaf 2012, Glasser et al. 2014, Shafee et al. 2015, Lifshits et al. 2018, Shamir et al. 2019). More specifically, a recently published methodology offers a framework for MRI-based quantification and visualization of the cortical laminar composition (Shamir et al. 2019).

By integrating the laminar structure derived from this framework into connectomics, not only we will be able to give a more detailed and comprehensive representation of brain connectivity, but we will also open a new avenue to future prospects in the field of connectomics. Integrating laminar composition into structural connectivity in this study enables us to introduce laminar connectivity, similarly to the way in which integrating structural and functional connectivity has led to effective connectivity. In the future, laminar and effective connectivity could conceivably be integrated into effective laminar connectivity.

In this study, we revisit Felleman and Van Essen’s 1991 work and attempt to recreate it on 7 excised macaque brains, using MRI techniques to model and investigate the laminar connectivity of its cerebral cortex. We do so while considering both the laminar organization as well as the interconnections of the cortex. We hypothesize that MRI, or more specifically diffusion weighted imaging (DWI), inversion recovery (IR) and T1w, can be integrated into a whole-brain laminar connectome through a simple granularity-based model based on published histological findings (Shamir and Assaf 2021).

This model is based on a systematic review of 51 prominent publications regarding laminar connectivity, mainly using histological and tract tracing methods. The model partitions the cortex into three laminar groups-supragranular (including layers 1-3), granular (layer 4) and infragranular (layers 5-6). It offers a simplified approach to overcoming the limitations of MRI connectomics by integrating whole-brain white and grey matter MRI datasets into a laminar-level connectome. The integration is completed using a set of laminar-level connectivity rules that are applied mainly on DWI white matter tractography, according to the locations and granularity indices of the connecting regions and weighted according to their cortical laminar compositions. A few additional rules are applied on assumed connections that are beyond the resolution of tractography.

The resulting whole-brain network of cortical laminar connectivity is then validated in the visual cortex by comparison to Felleman and Van Essen’s 1991 results. By using a more comprehensive definition of the cortex that takes into account its heterogenous laminar composition, we are able to use MRI neuroimaging to explore laminar-level structural connectomics.

## Methods and materials

### Macaque brains

Following are the steps taken for acquiring and processing the MRI images, resulting in the datasets of cortical grey matter substructure and white matter connectivity that are used as input for modelling and exploring the macaque laminar connectome.

### Sample

7 macaque brains were obtained from the Mammalian MRI (MaMI) database (Assaf et al. 2020). No animals were deliberately euthanized for the present study. Each brain was formalin fixated and some 24 hr before MRI, it was placed in phosphate-buffered saline for rehydration. For the MRI scan, each brain was placed in a plastic bag and immersed in fluorinated oil (Flourinert, 3 M). This procedure was done to minimize image artifacts caused by magnet susceptibility effects. Of the 7 macaque brains, one exemplary excised macaque brain was chosen for an expanded series of MRI scan protocols.

### MRI Acquisition

The excised macaque brain was scanned on a 7T/30 Bruker scanner with a 660 mT/m gradient system. The protocol was approved by the Tel Aviv University ethics committee on animal research. Three MRI sequences were performed:

1. DWI (diffusion weighted imaging) was performed with 4 segments diffusion weighted EPI sequence with a voxel size 0.48×0.48×0.48 mm3, image size 128×160×116 voxels, Δ/δ=20/3.3 ms, b=5000 s/mm2, with 96 gradient directions and additional 4 with b=0. The acquisition time for the DWI dataset was approximately 14 hr and 38 min. Performed for all macaque brains. DWI dataset was used for global white matter connectivity (tractography), using constrained spherical deconvolution (CSD) in ExploreDTI (Leemans et al. 2009).
2. T1w sequence was performed with a 3D modified driven equilibrium Fourier transform (MDEFT) sequence with: voxel size 0.2×0.2×0.2 mm3, image size 300×360×220 voxels, TR/TE=1300/2.9 ms, TI=400 ms. The acquisition time for the T1w dataset was approximately 2 hr and 13 min. Performed for the chosen exemplary macaque brain. This sequence was used as an anatomical reference with high gray/white matter contrast for segmentation and estimation of cortical surfaces (similar to clinical MPRAGE and SPGR sequences).
3. Inversion recovery was performed using a 3D FLASH sequence with: voxel size 0.67×0.67×0.67 mm3, image size 96×96×68 voxels, TR/TE=1300/4.672 ms and 44 inversion times spread between 25 ms up to 1,000 ms, each voxel fitted with up to 8 T1 values (similarly to Lifshits et al. 2018). The acquisition time for the IR dataset was approximately 69 hr and 34 min. Performed for the chosen exemplary macaque brain. T1w and inversion recovery datasets were used for cortical laminar composition analysis (similarly to the framework presented in Shamir et al. 2019).

### Cortical atlas and template

In order to conduct a comparison of our resulting model of cortical laminar connectivity in the macaque brain to the results presented by Felleman and Van Essen (Felleman and Van Essen 1991), we used their FVE91 atlas and FVE91 template (see Supplementary Material figure 1).

### Image registration

In the interest of transferring all datasets to a single anatomical space, all the following were registered to the T1w image: T1 layer probability maps, DTI images, atlas and template. Registration was completed in SPM12 using a rigid body transformation for the MRI datasets (with the first image of each dataset as the source image) and normalization followed by registration for the atlas and template.

### Image processing

The IR dataset was used for grey matter substructure composition analysis, according to the following steps:

1. Cortical volume sampling: With the intention of providing accurate volumetric sampling of the complex geometry of the cortex, we use a novel framework for cortical laminar composition analysis (Shamir et al. 2019). This framework utilizes a sampling system of virtual cortical spheres with the intention of offering a simple solution for cortical curvature heterogeneity and partial volume effects.
2. IR decay function fit: The IR 3D FLASH data were used for multiple T1 analysis, by calculating T1 values and their corresponding partial volumes on a voxel-by-voxel basis. The IR datasets were fitted to the conventional inversion recovery decay function with up to 8 possible T1 components per voxel (Lifshits et al. 2018).
3. T1 probabilistic classification: The multiple T1 components were used for accurate whole-brain classification to brain tissues on a voxel-by-voxel basis. A T1 histogram representing all T1 values was then fitted to a probabilistic mixture model (similarly to the method shown in Lifshits et al. 2018, Barazany et al. 2012, and Shamir et al. 2019), consisting of 11 t-distributions corresponding to different types of brain tissue, including cortical layer components (Peel et al. 2000, Shamir et al. 2019).
4. Cortical volume sampling and composition analysis: Because of the resolution difference between the spherical sampling system (average radius of ∼0.5 mm) and the T1 data (0.67^3^ mm^3^), a sampling solution was implemented using subvoxel information (similarly to Shamir et al. 2019). The solution starts with partitioning of each voxel into subvoxels, followed by assignment of spherical volume weights and concludes with cortical composition analysis inside all cortical spheres.
5. DTI analysis and tractography: The DWI dataset was used for global white matter connectivity analysis via standard DTI analysis followed by constrained spherical deconvolution (Tournier 2007), implemented in ExploreDTI (Leemans et al. 2009).

For more details on each of the above-mentioned image processing steps see Supplementary Material.

### Cortical granularity indices

An atlas of cortical granularity indices was to assist in the subsequent modelling process, based on a map of cytoarchitectonic features across the primate cortex (Beul and Hilgetag 2019) (see Supplementary Material figure 5).

### Modelling cortical laminar connectivity

The resulting datasets of cortical grey matter substructure and white matter connectivity were used as input for modelling and exploring the macaque laminar connectome. Datasets were integrated using an MRI-based, data-driven model of cortical laminar connectivity (Shamir and Assaf 2021). The model is based on a systematic review of a sample of articles focused on laminar connectivity, offering a way to overcome specificity and resolution limitations in MRI connectomics and presenting a simplified approach to integrating whole-brain white and grey matter MRI datasets into laminar-level connectivity (for a summary of our pipeline for modelling cortical laminar connectivity see figure 1).

**Fig. 1.**
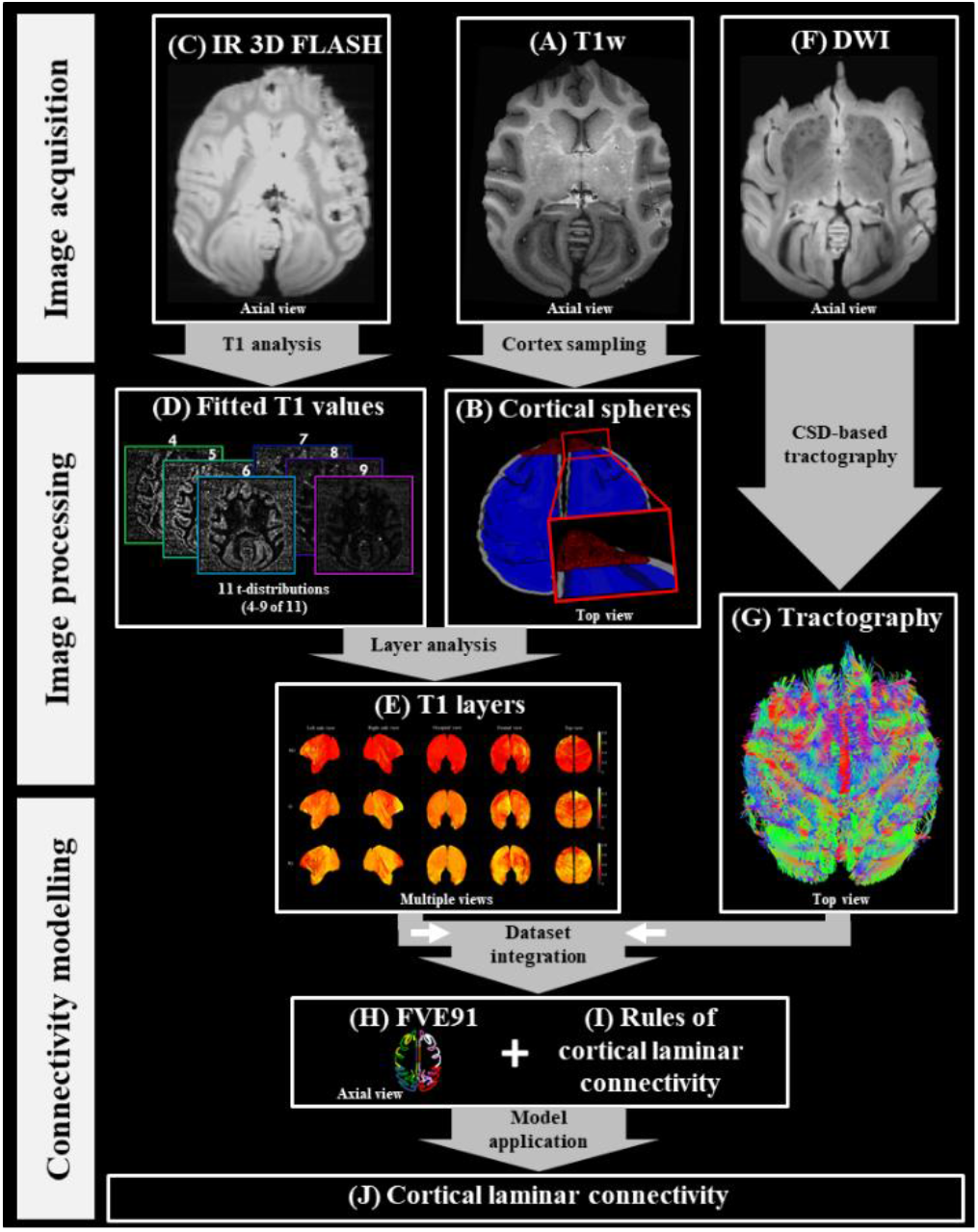
A streamline summary of our proposed pipeline for modelling cortical laminar connectivity, including the following steps: A- T1w image acquisition: for anatomical reference B- Cortical spheres: sampling volume between delineated cortical surfaces C- IR 3D FLASH image acquisition: for cortical laminar composition analysis D- Fitted T1 values: T1 analysis using a mixture model for tissue classification E- T1 layers: whole-brain laminar composition analyzed in spherical sampling system F- DWI image acquisition: for global white matter connectivity analysis G- Tractography: whole-brain tract analysis using constrained spherical deconvolution H- FVE91: both datasets analyzed in the Felleman and Van Essen atlas space I- Rules of cortical laminar connectivity: for integrating grey and white matter datasets J- Cortical laminar connectivity: the resulting whole-brain laminar-level connectome

### Human subject

The entire framework for cortical laminar connectivity was also applied on a single exemplary human subject. The subject was neurologically and radiologically healthy with no history of neurological diseases. The subject signed informed consent before enrollment in the study. The imaging protocol was approved by the institutional review boards of Sheba Medical Center and Tel Aviv University, where the MRI investigations were performed.

The subject was scanned on a 3T Magnetom Siemens Prisma (Siemens, Erlangen, Germany) scanner with a 64-channel RF coil. One MRI sequence was used to map the cortical connectome (1) and two T1-weighted MRI sequences were used to characterize the cortical layers (2, 3):

1. A standard diffusion-weighted imaging (DWI) sequence, with the following parameters: Δ/δ=60/15.5 ms, b max=5000 (0 250 1000 3000 & 5000) s/mm2, with 87 gradient directions, FoV 204 mm, maxG= 7.2, TR=5200 ms, TE=118 ms, voxel size 1.5×1.5×1.5 mm^3^, image size 128×128×94 voxels. This sequence was used for global white matter connectivity analysis.
2. An MPRAGE sequence, with the following parameters: TR/TE = 1750/2.6 ms, TI = 900 ms, voxel size 1×1×1 mm^3^, image size 224×224×160 voxels, each voxel fitted with a single T1 value. This sequence was used as an anatomical reference with high gray/white matter contrast.
3. An inversion recovery echo planar imaging (IR EPI) sequence, with the following parameters: TR/TE = 10,000/30 ms and 60 inversion times spread between 50 ms up to 3,000 ms, voxel size 3×3×3 *mm*^3^, image size 68×68×42 voxels, each voxel fitted with up to 7 T1 values (Lifshits et al. 2018). The acquisition time for the inversion recovery data set was approximately 12 min. This sequence was used for cortical laminar composition analysis.

For more details on each of the above-mentioned image processing steps for the human subject, see Supplementary Material: Model application on human subject).

## Results

In this study we aim to recreate Felleman and Van Essen’s 1991 work noninvasively using multimodal magnetic imaging of the macaque brain, including both its grey matter laminar structure and its white matter cortical connectivity. These datasets are then integrated using our model of cortical laminar connectivity and the resulting connectome is then further explored on the laminar level.

We will present first the whole-brain grey matter composition analysis, followed by white matter connectivity analysis: whole-brain structural connectome as well as V1 comparison to Felleman and Van Essen’s 1991 work. Then we will present the model of cortical laminar connectivity, validated by comparison to Felleman and Van Essen’s results, and close with complex network analysis of the resulting network.

### T1 layer composition analysis

Following image processing steps 1-4, T1 layers were estimated across the cortical volume. Granular layer 4 (T1 layer 4) was identified according to high probability in V1 and low probability in M1. Since the more myelinated layers lie deeper in the cortex, T1 layers were then easily categorized as supragranular (SG), granular (G) and infragranular (IG) and presented as surface overlays (see figure 2).

**Fig. 2.**
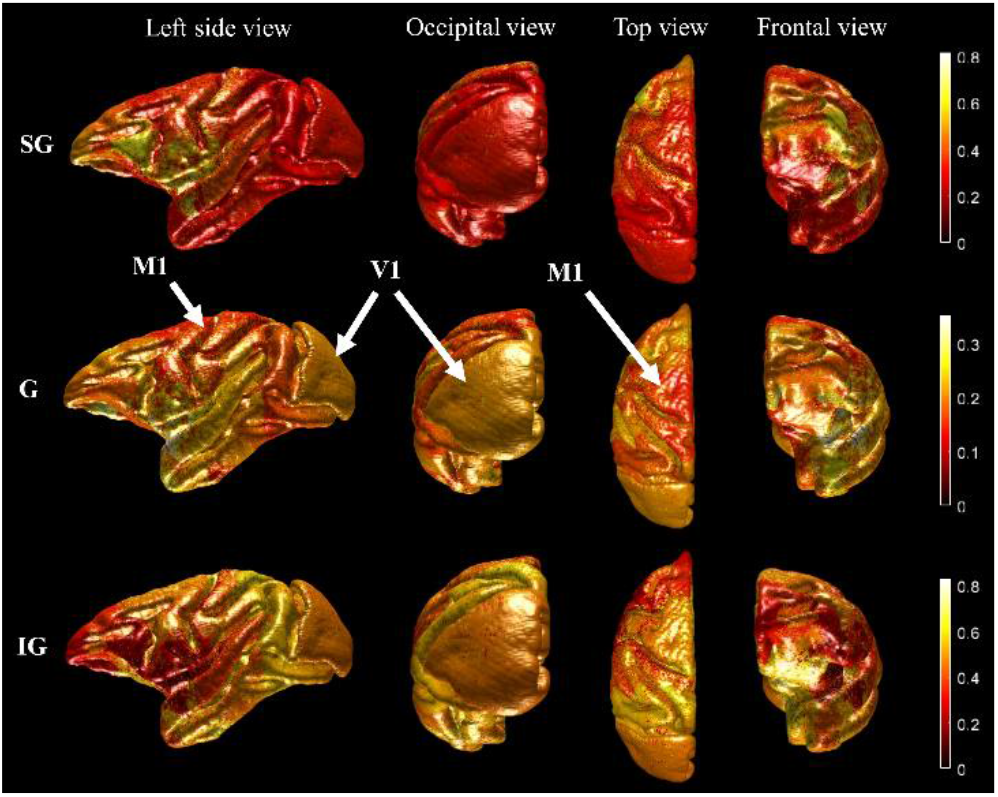
Macaque cortical laminar probability maps across the left hemisphere: Top row: supragranular (SG) composition, corresponding to T1 layers 1-3; Mid row: granular (G) composition, corresponding to T1 layer 4; Bottom row: infragranular (IG) composition, corresponding to T1 layers 5-6; Laminar components are scaled according to ranges of minimal to maximal values; Arrows indicate the notable features: high granular presence in the primary visual cortex (V1) and low granular presence in the primary motor cortex (M1)

### Standard cortical connectivity

Whole brain DTI tractography was completed using CSD (constrained spherical deconvolution) across all FVE91 regions (see Supplementary Material figure 4). For a full visualization of all model input datasets see figure 3, part 1 (below).

**Fig. 3.**
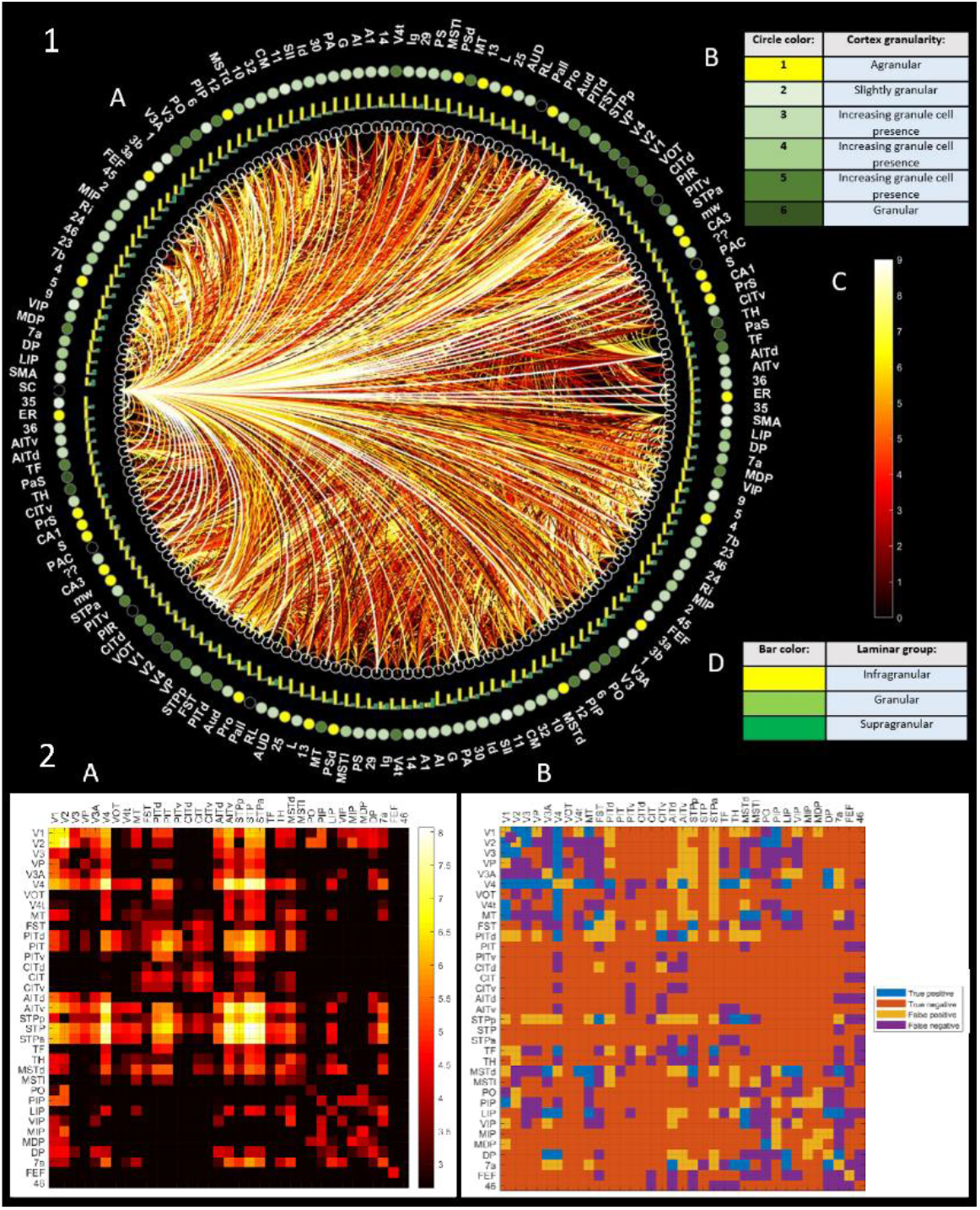
Macaque cortical connectivity: **Part 1 (top):** A circular network graph representing all model input datasets for the exemplary macaque brain (A), including: B- Legend for granularity indices, located in the outermost circle; C- Color scale for global white matter connectomics, representing connection strength of edges; D- Legend for color bars, representing grey matter laminar composition, located in the first outer circle; (Visualized using Circular-Connectome toolbox, available at: github.com/ittais/Circular-Connectome) **Part 2 (bottom):** Macaque visual cortex connectivity: A- Connectivity matrix, representing log(number of tracts) in visual regions (from tractography) B- Matrix representing comparison of connectivity resulting from the two different methods: predicted values (from tractography- A) to actual results (from tracing study- B)

DTI tractography in the visual cortex was compared to Felleman and Van Essen’s reported connections between 35 regions of the visual cortex (see Supplementary Material figure 6). Comparison of connectivity matrices demonstrates comparable general patterns of connectivity, with specific regions with higher interconnectedness (V1, V2, V3, Vp, V3A, V4) and others with very sparse interconnectedness (such as CIT and TH). For a dendrogram plot of the hierarchical relationship between clustered regions in the macaque visual cortex, see Supplementary Material figure 7.

In order to quantitatively compare the results in the visual cortex, we binarized both connectivity matrices. In the tractography-based connectivity matrix, existing connections were marked as ‘1’ and missing connections as ‘0’. In the connectivity matrix adapted from Felleman and Van Essesn’s study, connections categorized as ‘reported and published’ and ‘identified but unpublished’ were marked as ‘1’, while those categorized as ‘not reported’ and ‘conflicting reports’ were marked as ‘0’. We used Felleman and Van Essen’s reported connectivity as the actual results, and the tractography-based connectivity as the predicted results (see Supplementary Material figure 8).

Nine different routines for DWI dataset analysis were performed in order to evaluate their correspondence with Felleman and Van Essen’s cortical connectivity findings. All routines were performed in ExploreDTI (Leemans et al. 2009), using either CSD or DTI as the primary analysis method, with varying minimal tract lengths and/or seed fractional anisotropy (FA) values (see table 1). The results showcase the delicate balance between high sensitivity (true positive rate) and specificity (true negative rate) concerning tract identification in the macaque visual cortex. Higher sensitivity comes at a cost of lower specificity, and vice versa. Additionally, three classification values were evaluated: accuracy, which represents the rate of true classifications (both positive and negative), F1 score, which represents the harmonic mean of precision and recall, and positive predictive value, which represents the proportion of positive results.

**Table 1.** Evaluation of different fiber tracking routines in the macaque visual cortex: DWI dataset was analyzed using 9 different routines in ExploreDTI (Leemans et al. 2009), using either CSD or DTI as the primary analysis method, with varying minimal tract lengths and/or seed fractional anisotropy (FA) values; For each routine, the resulting cortical connectivity patterns were compared to Felleman and Van Essen’s 1991 findings in the visual cortex; Each comparison includes the following confusion matrix values: true positive rate (TPR), true negative rate (TNR), false positive rate (FPR), false negative rate (FNR), accuracy, F1 score and positive predictive value (PPV)

| # | Tractography method | Parameters | Cortical connectivity (tractography) |  |  |  |  |  |  |
| --- | --- | --- | --- | --- | --- | --- | --- | --- | --- |
|  |  |  | TPR | TNR | FPR | FNR | Accuracy | F1 score | PPV |
| 1 | CSD | Min L=5 | 46.5% | 62% | 38% | 53.5% | 58% | 0.348 | 0.278 |
| 2 | CSD | Min L=10 | 46% | 64% | 36% | 54% | 60% | 0.352 | 0.285 |
| 3 | CSD | Min L=20 | 42% | 66.5% | 33.5% | 58% | 61% | 0.336 | 0.28 |
| 4 | DTI | Min L=5 | 36% | 70.5% | 29.5% | 64% | 62% | 0.314 | 0.278 |
| 5 | DTI | Min L=10 | 35% | 71% | 29% | 65% | 63% | 0.309 | 0.278 |
| 6 | DTI | Min L=20 | 29% | 75% | 25% | 71% | 64% | 0.277 | 0.265 |
| 7 | DTI | Min L=5, FA=15 | 15.5% | 87.5% | 12.5% | 84.5% | 70% | 0.198 | 0.276 |
| 8 | DTI | Min L=10, FA=15 | 14% | 90% | 10% | 86% | 72% | 0.191 | 0.299 |
| 9 | DTI | Min L=20, FA=15 | 8% | 95% | 5% | 92% | 74% | 0.129 | 0.333 |

The first routine (CSD with min tract length of 5 mm) was chosen as our model input because it has the highest sensitivity value and the lowest false negative rate. Its relatively low accuracy value can be attributed to the low specificity, a fact apparent in its relatively high F1 score. We then estimated confusion matrix values across all connections, stating either true positive, true negative, false positive or false negative per connection (see figure 3, part 2). The resulting values: true positive rate- 46.5%, true negative rate- 62%, false positive rate- 38%, and false negative rate- 53.5%. In other words, the results exhibit relatively low sensitivity (46.5%) and relatively higher specificity (62%). The overall accuracy level, calculated as the sum of true positives and true negatives divided by sum of all positives and negatives, was estimated at ∼58%.

Once the routine was selected, we used a threshold to remove low connectivity values. The threshold was selected by evaluating positive predictive values (PPVs) for different threshold values across macaque brains (see Supplementary Material table 1), followed by averaging of the best performing values (resulting in a threshold of 13.5 tracts).

### Cortical laminar connectivity

### Whole-brain cortical laminar connectivity

In this section, we fuse the fiber tracking data (presented figure 3) with the cortical layer-granularity analysis results (presented in figure 2) in order to estimate the laminar connectivity properties. We used the Circular-Connectome toolbox (available at: github.com/ittais/Circular-Connectome) in order to display and explore the resulting model of whole-brain laminar connectivity (best viewed online in 3D).

For a visualization of the multilayered circular network graph representing the laminar connectivity of the macaque brain, see Supplementary Material figure 9.

### Visual cortex comparison of to Felleman and Van Essen findings

The resulting model of cortical laminar connectivity in the visual cortex was compared to an adaptation of Felleman and Van Essen’s reported laminar connections (see figure 4 below). The reported connections include origins and terminations of connections across 35 regions of the macaque visual cortex (based on Felleman and Van Essen 1991), summarized and adapted for comparison according to the following categories:

**Fig. 4.**
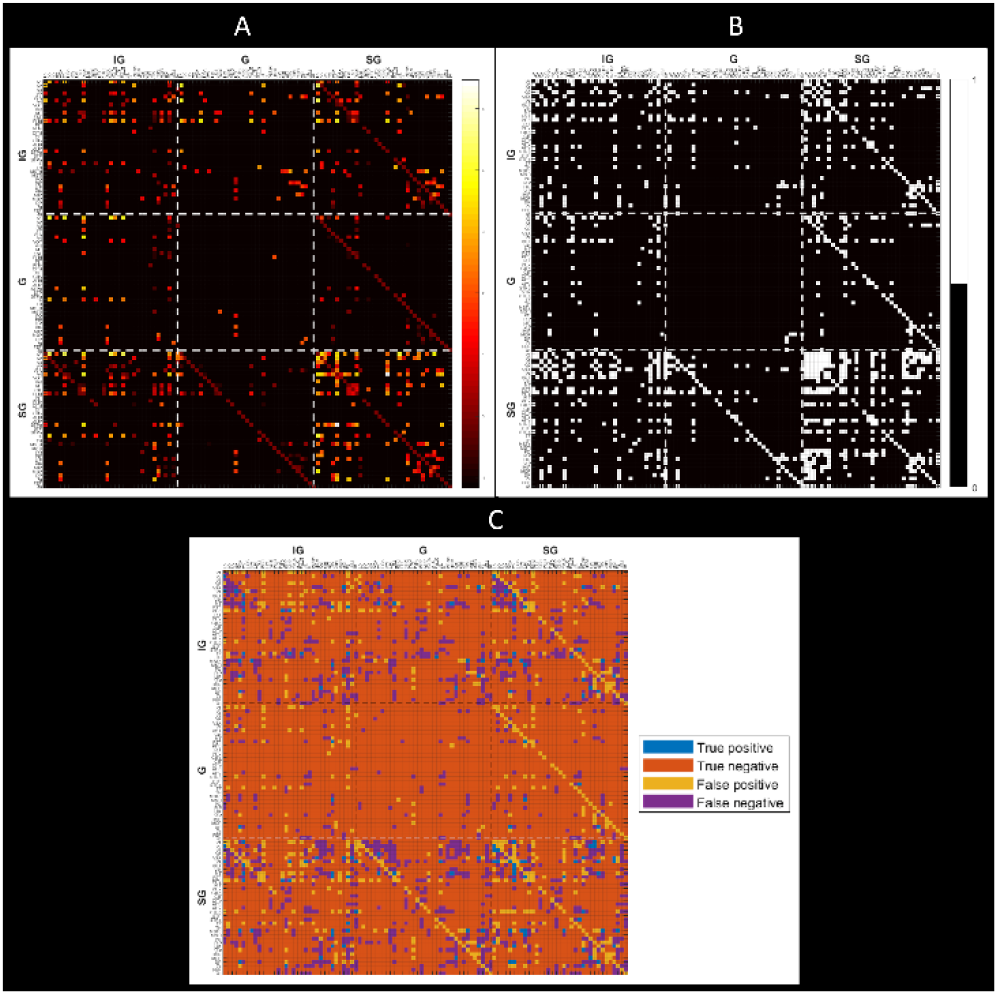
Macaque visual cortex laminar connectivity: A- Supra-adjacency matrix, representing laminar-level connections in visual cortex (model results, displayed as log(number of tracts)) B- Supra-adjacency matrix of reported cortical laminar connections, a summarized adaptation of connections reported in (Felleman and Van Essen 1991) C- Supra-adjacency matrix representing comparison of predicted values (from model) to actual results (adapted reports) All three matrices include infra-granular (IG), granular (G) and supra-granular (SG) origins and terminations of connections across 35 regions of the macaque visual cortex

a. Laminar location of origin/termination (as reported by Felleman and Van Essen 1991) on left, and the equivalent laminar group in our model on right:
  1. Origin: supralaminar (S)- SG, bilaminar (B)- IG+SG, infragranular (I)- IG;
  2. Termination: layer 4 (F)- G, columnar (C)- IG+G+SG, multilayer avoiding layer 4 (M)- IG+SG Where: IG- infragranular, G- granular (G), SG- supragranular.
b. If pathway features are reported partially- all laminar categories were marked.

The adaptation results in a supra-adjacency matrix summary of reported cortical laminar connections across the macaque visual cortex (see figure 4 B).

Visual comparison of adjacency matrices demonstrates similarities in unique patterns of connectivity, including generally higher connectedness of infragranular and supragranular laminar compartments, relative to granular connectedness. More specifically, the comparison demonstrates high interconnectedness of supragranular to supragranular compartments and sparse interconnectedness of granular to granular compartments. One visual way in which the results of the model differ from Felleman and Van Essen’s is their diagonal connections, as a result of intra-regional vertical connections in the model. These connections are not tractography-based, but rather assumed connections as part of an adaptation of the canonical circuit (Shamir and Assaf 2021). Despite their visual presence, these intraregional connections represent a small portion of connections (about 0.3% of all possible connections), and therefore have a marginal effect on the overall model accuracy.

In order to quantitatively compare the results in the visual cortex, once again we binarized both laminar connectivity matrices (see Supplementary Material figure 10). Similarly to the method presented earlier, we used Felleman and Van Essen’s reported laminar connectivity as the actual results, and the resulting model of cortical laminar connectivity as the predicted results.

The nine different routines for DWI dataset analysis were also used in order to evaluate the each resulting model’s correspondence with Felleman and Van Essen’s cortical laminar connectivity findings (see table 2). The balance between sensitivity and specificity appears here as well, where the first routine (CSD with min tract length of 5 mm) exhibits the highest sensitivity and a relatively high F1 score.

**Table 2.** Evaluation of the effect of different fiber tracking routines on the resulting model of laminar cortical connectivity in the macaque visual cortex: DWI dataset was analyzed using 9 different routines in ExploreDTI (Leemans et al. 2009), using either CSD or DTI as the primary analysis method, with varying minimal tract lengths and/or seed fractional anisotropy (FA) values (same as table 1); For each routine, the resulting patterns of laminar connectivity were compared to Felleman and Van Essen’s 1991 findings. For each comparison, the following confusion matrix values were evaluated: true positive rate (TPR), true negative rate (TNR), false positive rate (FPR), false negative rate (FNR), accuracy and F1 score

| # | Tractography method | Parameters | Cortical laminar connectivity (model) |  |  |  |  |  |
| --- | --- | --- | --- | --- | --- | --- | --- | --- |
|  |  |  | TPR | TNR | FPR | FNR | Accuracy | F1 score |
| 1 | CSD | Min L=5 | 22% | 86% | 14% | 78% | 78% | 0.198 |
| 2 | CSD | Min L=10 | 21.5% | 86.5% | 13.5% | 78.5% | 79% | **0.199** |
| 3 | CSD | Min L=20 | 18% | 88% | 12% | 82% | 79% | 0.177 |
| 4 | DTI | Min L=5 | 14.5% | 89.5% | 10.5% | 85.5% | 80% | 0.155 |
| 5 | DTI | Min L=10 | 14.5% | 90% | 10% | 85.5% | 81% | 0.156 |
| 6 | DTI | Min L=20 | 11% | 91% | 9% | 89% | 81% | 0.127 |
| 7 | DTI | Min L=5, FA=15 | 7% | 94% | 6% | 93% | 83% | 0.0944 |
| 8 | DTI | Min L=10, FA=15 | 6% | 95% | 5% | 94% | 84% | 0.086 |
| 9 | DTI | Min L=20, FA=15 | 3% | **96.5%** | **3.5%** | 97% | **85%** | 0.048 |

Once again, the first routine (CSD with min tract length of 5 mm) exhibited the highest sensitivity value and the lowest false negative rate. We then estimated confusion matrix values for this routine across all connections, stating either true positive, true negative, false positive or false negative per connection (see figure 4 C). The resulting values: true positive rate- 22%, true negative rate- 86%, false positive rate- 14%, and false negative rate- 78%. The results exhibit low sensitivity (22%, highest of all routines), high specificity (86%), and relatively high accuracy (78%).

The connectivity analysis process was repeated for 6 additional excised macaque brains, all from the Mammalian MRI (MaMI) database (Assaf et al. 2020). The T1 layer composition of the original macaque brain was used for all the additional 6 macaque brains. The connectivity analysis includes standard cortical connectivity and cortical laminar connectivity modelling (for an analysis of consistency in connectivity patterns in the visual cortex across all macaque brains, see Supplementary Material figure 11). Accuracy values were evaluated for all macaque brains, including both standard cortical connectivity and modelled cortical laminar connectivity (see Supplementary Material table 2). The evaluation of all 7 macaques reveals relatively low standard deviation values, thus demonstrating a robustness of our modelling framework.

Patterns of cortical and cortical laminar connectivity were averaged for all 7 macaque brains. In order to compare the resulting connectivity patterns in the visual to Felleman and Van Essen’s findings, once again we binarized the connectivity matrices. Since the resulting average cortical and cortical laminar connectivity matrices are dense, we used the median value of each as thresholds for binarization (see figure 5). Consequently, for the average macaque brain the accuracy value was estimated at ∼63% for cortical connectivity, and at ∼79% for cortical laminar connectivity.

**Fig. 5.**
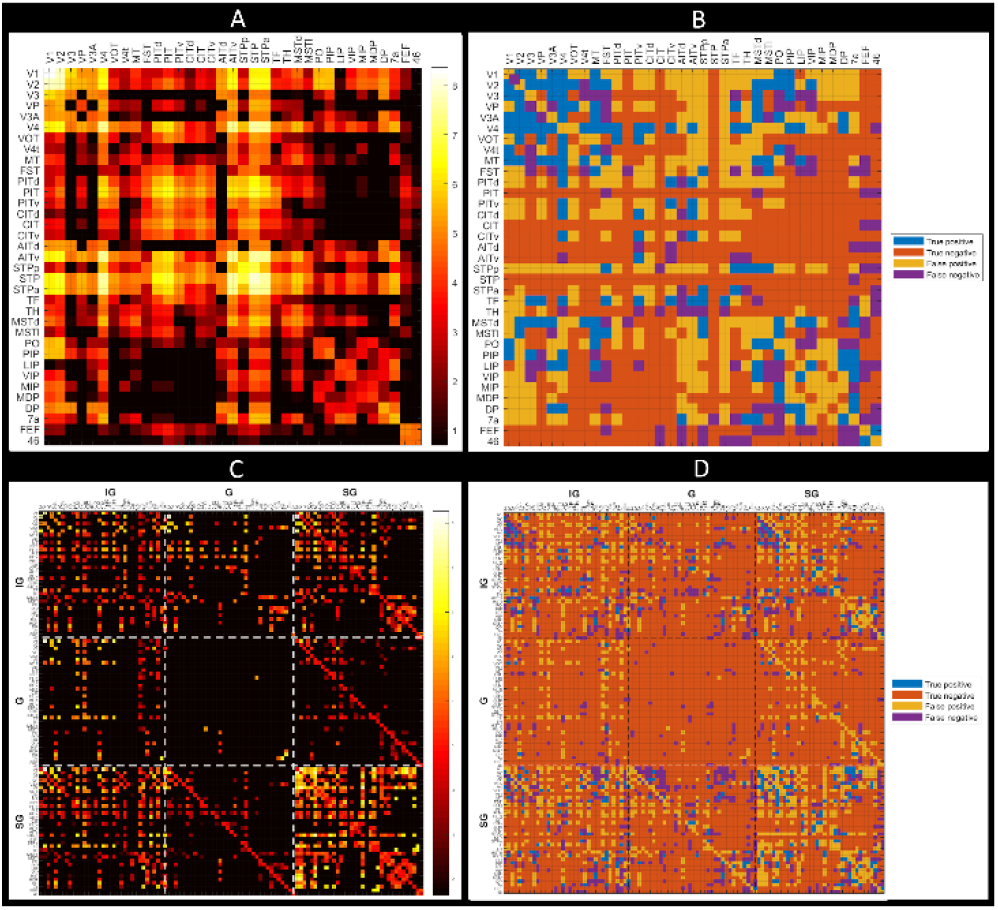
Average cortical connectivity (top) and cortical laminar connectivity (bottom) in the visual cortex for 7 macaque brains: A- Supra-adjacency matrix, representing laminar-level connections in visual cortex (model results, displayed as log(number of tracts)) B- Supra-adjacency matrix representing comparison of predicted values (from model) to actual results (adapted reports) C- Supra-adjacency matrix, representing laminar-level connections in visual cortex (model results, displayed as log(number of tracts)) D- Supra-adjacency matrix representing comparison of predicted values (from model) to actual results (adapted reports)

### Complex network analysis

We explored the modelled laminar connectome using complex network analysis tools, similarly to the way in which standard connectomics are explored. In the past, criteria such as rich-club organization (Van den Heuvel and Sporns 2011) and connectivity hubs (Van den Heuvel and Sporns 2013) have been used to gain insight into the organization of the connectome. These criteria and others could easily be adapted and utilized to further explore the modelled laminar connectome.

We chose three straightforward node centrality criteria for exploring both the standard connectome as well as the laminar connectome of the macaque:

1. Node degree indicates the number of edges connected to that node.
2. Node betweenness indicates how often the node appears on a shortest path between two nodes in the network.
3. Nodes closeness indicates the reciprocal of the sum of the length of the shortest paths between the node and all other nodes.

The complex network analysis showcases differences not only between the standard connectome and the modelled laminar connectome, but also between the three components of the laminar connectome (see figure 6). Comparison of the degree criterion between the standard and laminar connectomes demonstrates the centrality of the subcortex in the standard connectome as well as the infragranular and supragranular components of the laminar connectome. It appears that the infra- and supragranular layers have a stronger overall contribution to connectedness in the laminar connectome, while in the granular component the occipital lobe and particularly the visual cortex exhibit relatively high centrality and higher degree values. High degree values can also be seen in the supragranular component in the frontal lobe around the motor cortex. Some interhemispheric asymmetry can be seen, specifically in the infragranular and granular components of the laminar connectome.

**Fig. 6.**
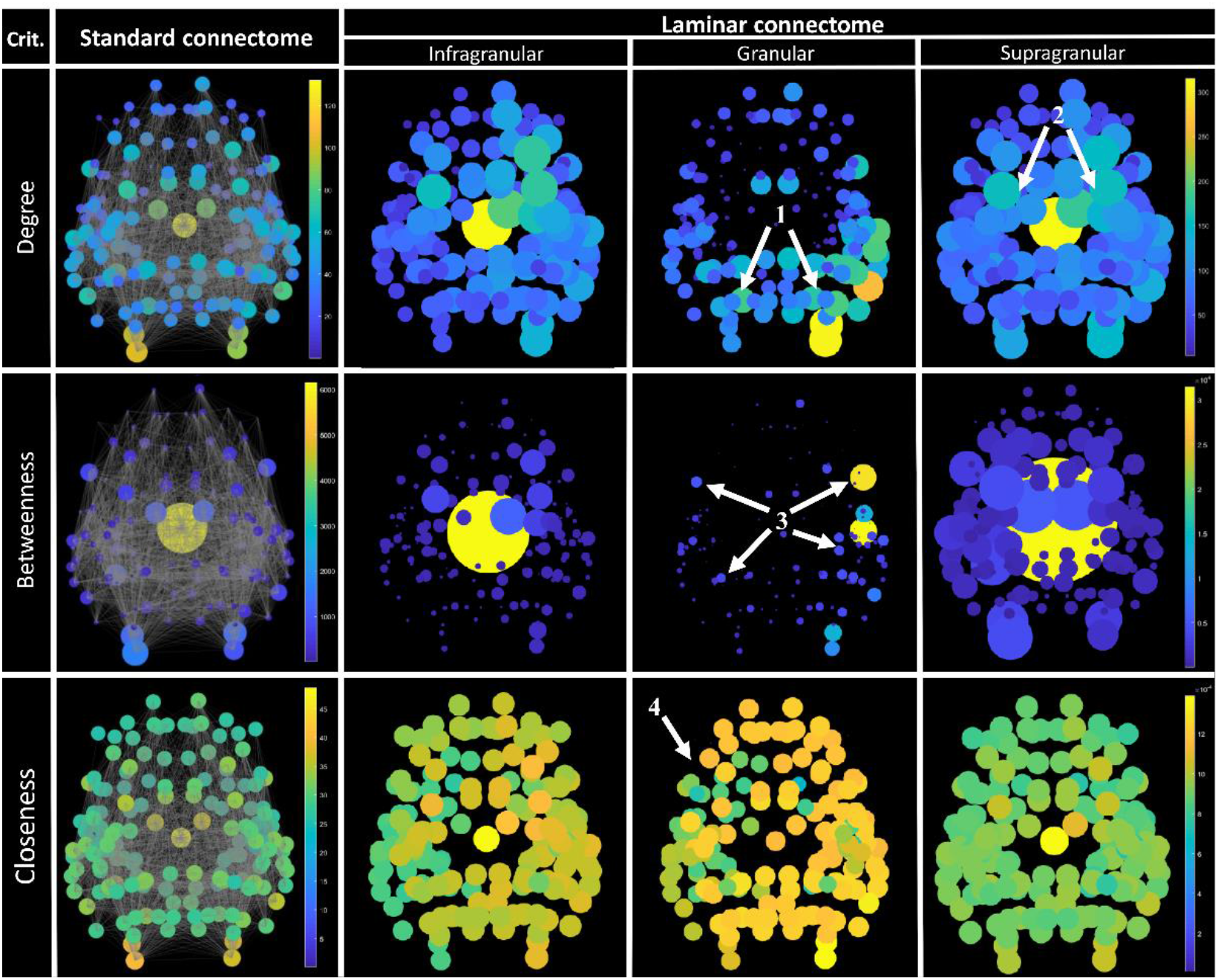
Comparison of network criteria of standard versus laminar macaque connectomes: Columns: first column represents standard connectome; second, third and fourth columns represent laminar connectome components (infragranular, granular and supragranular components respectively); Rows: top row includes node degrees, mid row includes node betweenness and bottom row includes node closeness (all represented across FVE91 regions); All images from a top view, with node values scaling both circle size and color (legend on right) Arrows indicate the notable features: 1- High degree values in the granular component in the visual cortex 2- High degree values in the supragranular component in the frontal lobe, around the motor cortex 3- ‘ 3- Hubs’ with high betweenness values in the granular component across both hemispheres 4- Higher closeness values overall in the granular components, relative to the other laminar components

Comparison of the betweenness criterion between the two connectomes demonstrates the increasing centrality of the subcortex, starting from the standard connectome and progressively in the infragranular and expressly the supragranular connectome. The granular component displays once again high centrality of the visual cortex. Several other betweenness ‘hubs’ in the granular component also appear in more frontal regions across both hemispheres. Some interhemispheric asymmetry can be seen across all connectomes and their components.

Comparison of the closeness criterion demonstrates relatively few regional differences within both types of connectome. Nonetheless, there are notable scale differences between the two, where the standard connectome exhibits significantly higher values compared to the laminar connectome. These differences can be explained by the larger number of nodes in the laminar connectome, compared to the standard connectome, resulting in a higher sum of shortest paths to all nodes and a lower resulting value. When comparing individual laminar network criteria, the granular layer demonstrates slightly higher closeness values across the entire cortex.

### Model application on human subject

The entire framework for modelling cortical laminar connectivity presented here was also applied on a single exemplary human subject across Von Economo-Koskinas atlas regions (for more on the application of the framework, see Supplementary Material: Model application on human subject). The model input and the resulting model of cortical laminar connectivity are both visualized in figure 7 (below).

**Fig. 7.**
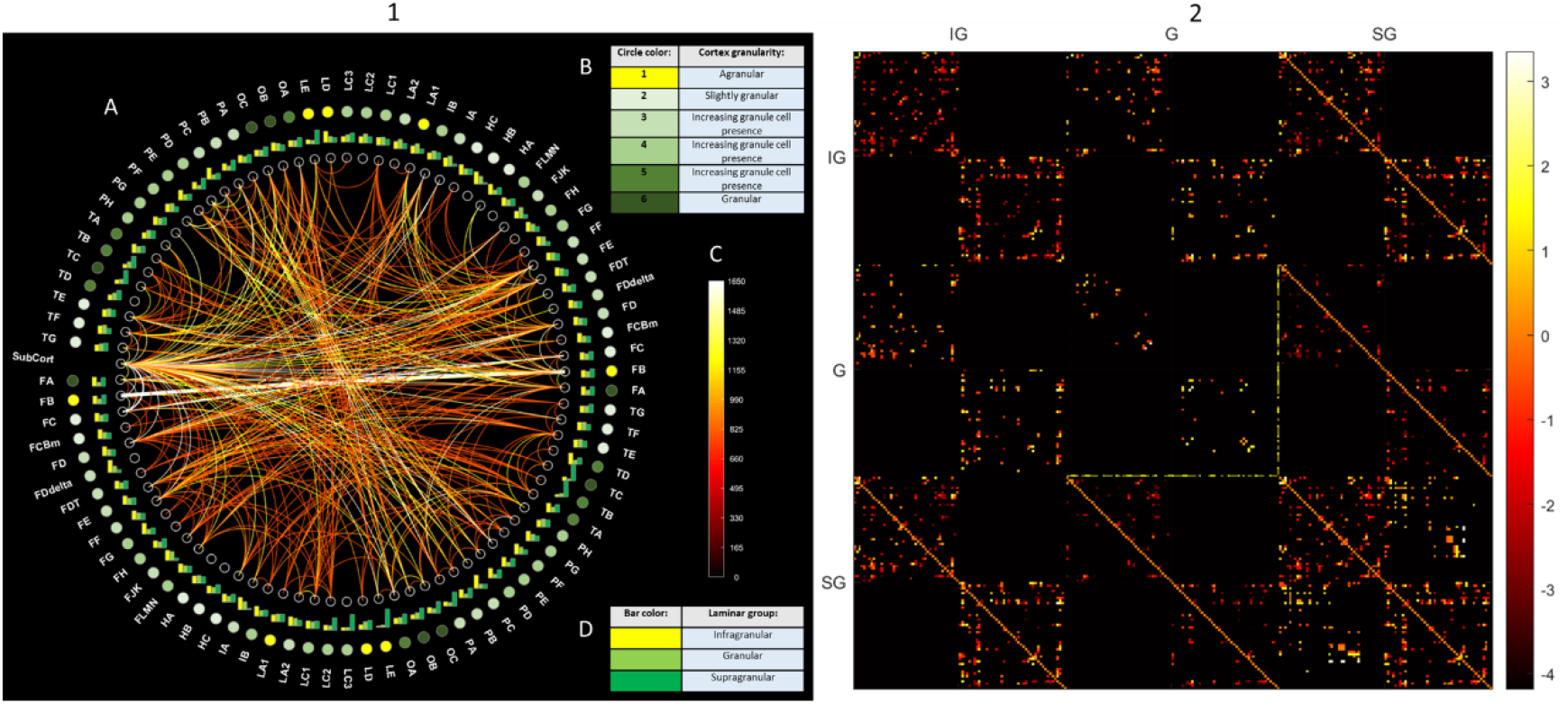
Modelling cortical laminar connectivity in the human: 1- Model input: A- a circular network graph of all model input datasets (A), including: B- Legend for granularity indices, located in the outermost circle; C- Color scale for global white matter connectomics, representing connection strength of edges; D- Legend for color bars, representing grey matter laminar composition, located in the first outer circle; (visualized using Circular-Connectome toolbox, available at: github.com/ittais/Circular-Connectome) 2- Model output: supra-adjacency matrix, representing whole-brain laminar-level connections across Von Economo- Koskinas atlas regions (model results, displayed as log(number of tracts)) Where: IG- infragranular, G- granular, SG- supragranular

For a comparison of network criteria of standard versus laminar connectomes for the human subject, see Supplementary Material figure 16.

## Discussion

In this study, we use multi-modal MRI techniques to model and explore the laminar connectome of the macaque brain. By using a data-driven model, we were able to combine results from diffusion-weighted imaging and T1-weighted imaging in order to estimate cortical laminar connectivity across the macaque brain. This innovative model into account not only the white matter interconnections of the cerebral cortex, but also its grey matter heterogenous laminar substructure.

The choice to focus on the macaque brain was motivated by a desire validate the results in the visual cortex by comparison to Felleman and Van Essen’s seminal study about hierarchical processing in the primate visual cortex (Felleman and Van Essen 1991). We recreated the macaque laminar connectivity in the visual cortex and across the entirety of the cortex non-invasively.

On the subject cortical grey matter composition, we used the unique framework for cortical sampling and analysis (Shamir et al. 2019). The implementing of this framework for the first time on a nonhuman subject was validated using granularity-based benchmarks, including high granular presence in the primary visual cortex and low granular presence in the primary motor cortex. Consequently, we gave further validation to the framework and to previous studies on cortical laminar analysis (Barazany and Assaf 2012, Lifshits et al. 2018).

After implementing and comparing several routines for standard connectivity analysis on the diffusion weighted imaging dataset, we chose first routine (CSD with min tract length of 5 mm) as our model input. It is worth noting that while this routine has the highest sensitivity and lowest false negative rate of all nine routines, its accuracy levels are still considered rather low (sensitivity- 46.5%, specificity- 62%, accuracy- 58%). This deficiency may be attributed to several factors, primarily relating to either DTI limitations or methodological issues. The limitations of DTI are highlighted when dealing with nonhuman excised and fixated specimens and further intensified when our ‘gold standard’ for comparison is a dataset based on histological tract-tracing, a vastly different methodology from MRI. In addition, it appears that even the best fiber tracking routine misses some tracts, as evident in the false negative rate- 53.5%. Concurrently, the best tracking routine still incorrectly reconstructs other tracts, as evident in numerous anatomically improbable short fibers close to the outer cortical surface (seen in Supplementary Material figure 4). With regards to methodological limitations, the most prominent limitation in this case relates to imperfect registration to small visual regions in the Felleman and Van Essen cortical atlas.

After selecting the best fitting routine for standard connectivity analysis, white and grey matter datasets were integrated through a novel model of cortical laminar connectivity (Shamir and Assaf 2021). We used the Circular-Connectome toolbox (available at: github.com/ittais/Circular-Connectome) to visualize both the model input datasets as well as the resulting model output in circular graph formats.

In an effort to validate the results, we compared the modelled laminar connectome in the visual cortex to Felleman and Van Essen’s widely reported 1991 study on histological tracing results. Their results were adapted and analyzed to match the model division of each regional node into three granularity-based laminar components. The comparison included 35 regions across the macaque visual cortex.

The comparison was conducted using both visual and quantitative assessment of the two adjacency matrices, showcasing reported and modelled laminar connections across infragranular, granular and supragranular laminar locations across 35 visual regions. Visual assessment showcased general similarities such as higher connectedness of infragranular and supragranular laminar compartments, relative to granular connectedness. Another similarity is exhibited in higher laminar interconnectedness within the supragranular laminar compartment, particularly compared to the sparse interconnectedness within the granular compartment. One visual aspect in which the results differ from Felleman and Van Essen’s is diagonal connections, because of intra-regional vertical connections in the model. It is well worth stating that these connections are assumed, rather than tractography-based. They are part of an adaptation of the canonical circuit, which can easily be tweaked and adjusted. Nonetheless, these connections represent a relatively small percentage of overall connections and consequently their effect on overall accuracy levels is peripheral.

Quantitative comparison of the two adjacency matrix showcased low sensitivity (22%), high specificity (86%), and relatively high accuracy (78%). The overall model accuracy in the visual cortex, as compared to Felleman and Van Essen’s work, was estimated at an average of about 83% for all seven macaque brains. The relatively low true positive rates and relatively high false negative rates can be attributed to similar patterns in cortical connectivity. These patterns in turn can be ascribed to limitations in DTI tractography even within the best evaluated fiber tracking routine, such as missed tracts and incorrectly reconstructed tracts (as discussed earlier). With regards to model limitations, specific model assumptions regarding assumed connections may negatively affect accuracy levels (notice in figure 4 C patterns of false negatives, like those in standard connectomics, and false positives across diagonals, respectively). An additional limiting factor is suboptimal tract tracing, due to dependency on the injection procedure and the selection of the injection site. These limitations can cause some cortical connections to become less or more apparent and subsequently affect the modelled laminar connections.

This study offers a notable application and validation of the proposed novel neuroimaging model of cortical laminar connectivity in the macaque visual cortex. The validity of the model is strengthened by the resulting average accuracy level, which was evaluated at 83%. Nonetheless, the proposed model does not solve the current limitations of DTI and tractography, as revealed in the limited accuracy levels of standard connectomics and consequently in the accuracy of the laminar connectome. What the model does offer is a robust framework for exploring cortical connectivity on the laminar level that is applicable today and will remain applicable in the future as the imaging and methodology for connectivity analysis advance. As the standard connectome becomes more accurate, so will the modelling framework result in a more precise delineation of the connectivity patterns on the laminar level.

Following this validation of the model of cortical laminar connectivity in the macaque brain, a second comparison conducted included the use of network analysis tools to compare the laminar connectome to the standard connectome on which it is based in part. Three centrality criteria were estimated and compared across connectome and laminar components: degree, betweenness and closeness.

Notable features can be seen specifically in the granular component of the laminar connectome, where high degree values appear in the visual cortex, ‘hubs with high betweenness values appear across both hemispheres, and overall higher closeness values appear relative to the other laminar components. Furthermore, the supragranular component showcases high degree values in the frontal lobe in general, and in the motor cortex in particular.

In the future, this modelling framework can be implemented on groups of human subjects to explore and characterize patterns of cortical connectivity on the laminar level. In addition to providing a more comprehensive description of the human connectome, our framework has the potential to help quantify finer aspects of connectivity that may reveal hidden mechanisms of brain integration in both health and pathologies. Findings in pathologies such as schizophrenia (Garey 2010, van Berlekom et al. 2020) and conditions such autism (Minshew and Williams 2007, Zikopoulos and Barbas 2013) have revealed abnormalities in cortical connectivity that are presumed to be layer dependent. Our laminar-level analysis of cortical connectivity could help elucidate the connectivity patterns behind such conditions.

## Information Sharing Statement

The source code of the present model connectomes (standard and laminar) and the codes for the circular graphs (expanded and multilayered, respectively) are freely available for non-commercial use from github.com/ittais/Circular-Connectome.

