## Supplementary Material for "Modelling cortical laminar connectivity in the macaque brain"

Ittai Shamir [a] and Yaniv Assaf [a,b]

[a] Department of Neurobiology, Faculty of Life Sciences, Tel Aviv University, Tel Aviv, Israel

[b] Sagol School of Neuroscience, Tel Aviv University, Tel Aviv, Israel

* Corresponding author:
Ittai Shamir
Address: Dept. of Neurobiology, Tel Aviv University, Ramat Aviv, Tel Aviv, 69978, Israel

ORDCID: 0000-0003-4028-5154

### Methods and materials

#### Cortical atlas and template

In order to conduct a comparison of our resulting model of cortical laminar connectivity in the macaque brain to the results presented by Felleman and Van Essen (Felleman and Van Essen 1991), we used the following cortical atlas and template (see figure 1):

1. Macaque cortical atlas (see figure 1 A): the FVE91 parcellation divides the cortex into 84 regions, as described by Daniel J. Felleman and David C. Van Essen (Felleman and Van Essen 1991).
2. Macaque cortical template (see figure 1 B): the F99 space is defined by a 0.5 mm MRI scan of an approximately 5-year-old male macaque monkey case F99UA1, provided by M. Logothetis and introduced by David C. Van Essen (Van Essen 2002).

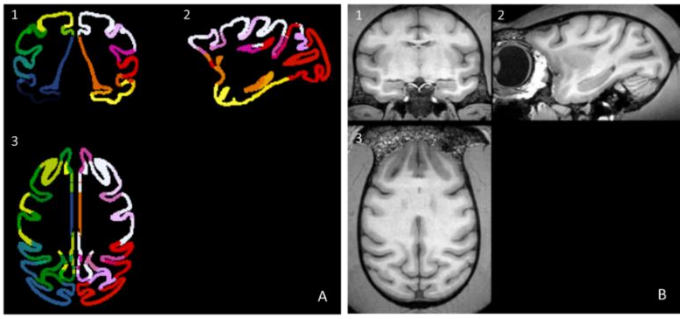


**Fig. 1** Macaque cortical atlas FVE91 (A) and template F99 space (B): Coronal (1), sagittal (2) and axial (3) views

#### Image processing

##### 1. Cortical volume sampling

In order to provide accurate volumetric sampling of the complex geometry of the cortex, we use the unique framework for cortical laminar composition analysis (presented by Shamir et al. 2019). This framework offers a simple solution for cortical curvature heterogeneity and the partial volume effects associated with cortical laminar analysis. Briefly, the framework uses a sampling system made up of virtual cortical spheres, as a symmetric and rotationally invariant volumetric alternative to cortical normal.

Since the subject is a non-human primate, we offer a straightforward alternative to the human MPRAGE analysis and segmentation of the *FreeSurfer* pipeline (Fischl 2012). The alternative utilizes the registered FVE91 macaque cortical atlas to delineate each cortical surface, using the steps:

a. Outer cortical surface: delineating the border between cortical GM bordering and the surrounding fluid:
   1. Binarize the macaque cortical atlas in order to create a mask
   2. Fill binarized atlas per cortical slice
   3. Rebuild entire filled hemispheric volumes from slices
   4. Split hemispheric volumes to left and right hemispheres in order to extract hemispheric surfaces
   5. Smooth both hemispheric volumes
   6. For each hemisphere, extract a triangular isosurface
   7. Resample each isosurface to ~150K vertices and ~300K faces

b. Inner cortical surface: delineating the border between cortical GM bordering and the underlying WM:
   1. Manually fill gaps in binarized cortical atlas (from step a. 1)
   2. Split cortical atlas to left and right hemispheres
   3. Per hemisphere, subtract cortical atlas (from step b. 2) from filled cortical volume (from step a.4)
   4. Smooth resulting binary images of hemispheric WM
   5. For each hemisphere, extract a triangular isosurface
   6. Resample each isosurface to ~150K vertices and ~300K faces in order to match original framework

c. Mid cortical surface: estimating the center of cortical GM:
   Run through all vertices of inner surface (~150K per hemisphere), for each:
   1. Calculate normal direction to inner surface (cortical normal)
   2. Find closest vertex on outer surface and calculate minimal distance (cortical width at that point)
   3. Estimate location of corresponding mid surface vertex:
   Mid surface vertex=Inner surface vertex+0.5*minimal distance (in normal direction)
   Once all mid surface vertices have been estimated, reconstruct entire mid triangular surface (see figure 2 A).

Once all three cortical surfaces were delineated per hemisphere, a system of virtual cortical spheres was built, using mid surface vertices and half the value of their corresponding cortical thickness as sphere centers and radii (respectively). In order to avoid excessive volumetric overlap of cortical spheres (Shamir et al. 2019), we subsampled the mid surface vertices by a factor of 2, to ~75,000 spheres per hemisphere (see figure 2 B).

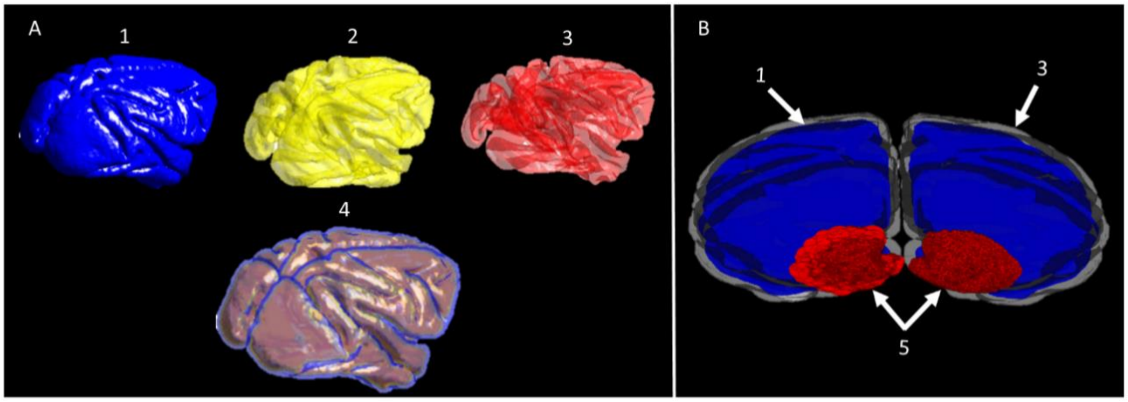


**Fig. 2** Macaque cortical volume sampling:

A- Right side view of delineated cortical surfaces: (1) outer, (2) mid and (3) inner cortical surfaces, and all three cortical surfaces (4). Cortical thickness values range from ~0.5 mm to ~3mm;

B- Occipital view of a small portion of the spherical sampling system, where the inner cortical surface is represented in opaque blue (1), the outer cortical surface is represented in clear gray (3) and the cortical spheres of the occipital lobes are represented in red (5)

#### 2. IR decay function fit

The IR 3D FLASH data were used for multiple T1 analysis, by calculating T1 values and their corresponding partial volumes on a voxel-by-voxel basis. The IR datasets were fitted to the conventional inversion recovery decay function with up to 8 possible T1 components per voxel (Lifshits et al. 2018):

$$\mathrm{M}(\mathrm{TI_i}) = \sum_{j=1}^{7} \mathrm{M0_j} \cdot \left(1 - 2\mathrm{e}^{-\mathrm{TI_i}/\mathrm{T_{1j}}}\right) \tag{1}$$

Where:

$\mathrm{M}(\mathrm{TI_i})$- Magnetization at the $i^{th}$ inversion recovery image, in other words the magnetization measured for each specific T1 component.

$\mathrm{M0_j}$- Predicted magnetization at TI=0ms for each T1 component (j) in the voxel.

$\mathrm{T_{1j}}$- Longitudinal relaxation time for each T1 component.

j was set up to 7 for the low-resolution experiments, indicating fit to seven individual exponential fits, based on the assumption that there are 8 T1 components in the tissue – 1 for CSF, 1 for WM and heavily myelinated layer of the cortex and additional 5 cortical layers.

Normalization of each of the predicted magnetization values according to $\frac{\mathrm{M0_j}}{\sum_{i=1}^{j} \mathrm{M0_j}}$ then represents the voxel contribution of each corresponding T1 component (j).

#### 3. T1 probabilistic classification

The multiple T1 components were used for accurate whole-brain classification to brain tissues on a voxel-by-voxel basis. A T1 histogram was initially plotted for this purpose. Each IR set of images consists of up to ~$5 \times 10^6$ potential T1 values: 96x96x68 [voxels] ×8 [T1 fitted values]. A single histogram was built, representing all fitted T1 values according to their partial volumes in each voxel across the image.

The T1 histogram was then fitted to a probabilistic mixture model (similarly to the method shown in Lifshits et al. 2018, Barazany et al. 2012, and Shamir et al. 2019), consisting of t-distributions (Peel et al. 2000, Shamir et al. 2019). The probability of each t-distribution in the voxel was calculated using Bayes' formula:

$$P_k = \sum_{i=1}^{8} f_i \cdot \frac{p\left(T_{1(i)}|k\right)p(k)}{p\left(T_{1(i)}\right)} \tag{2}$$

Where:

$k$- A specific t-distribution.

$T_{1(i)}$- T1-value of the $i^{th}$ component of the voxel.

$f_i$- Partial volume of $T_{1(i)}$ (normalized as show in previous section).

$p(T_1)$- General whole-brain probability of a $T_1$-value.

$p(k)$- Probability of t-distribution k.

$p(T_1|k)$- Probability of the $T_1$-value in t-distribution $k$.

This model was used as a means of probabilistic classification of T1 values into clusters representing different brain tissues. Evaluation of the appropriate number of t-distributions for the mixture model was completed using the Bayesian information criterion (BIC), a criterion used for probabilistic model selection. The mixture model was fitted repeatedly using a different number of distributions each time, ranging between 1-20 distributions, and evaluated using –log of BIC value.

Using this approach, fit to 11 t-distributions was deemed sufficient, where –log(BIC) reaches a maximal value with the fewest distributions. These 11 distinct T1 clusters correspond to different types of brain tissue (see figure 3):

1. White matter (WM): characterized by low T1 values, represented by t-distributions 1-3.
2. Gray matter (GM): characterized by mid-range T1 values, represented by $4^{th}$-$9^{th}$ t-distributions corresponding to 6 T1 layers, with decreasing degrees of myelination. Because of the link between myelination and shorter T1 values, T1 layer 4 ($4^{th}$ t-distribution) represents the most highly myelinated and innermost cortical component, with decreasing degrees of myelination moving outwards up to the outermost cortical component- T1 layer 1 ($9^{th}$ t-distribution).
3. Phosphate-buffered saline (PBS): excess fluid (no cerebral spinal fluid in an excised brain), characterized by high T1 values, represented by t-distributions 10-11 (also includes signal noise).

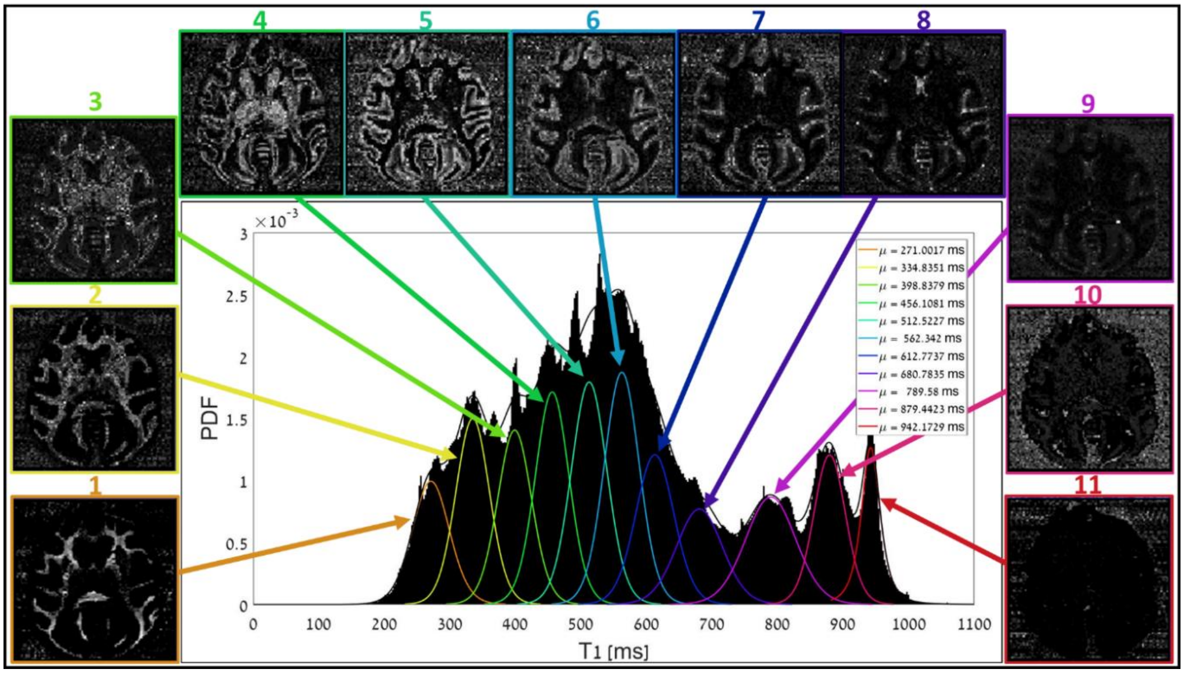


**Fig. 3** Cortical laminar composition analysis:
A histogram of T1 values (black) is fit to a mixture of 11 t-distributions (color map). Low T1 values represent white matter (1-3), high values represent fluid/noise (10-11), and mid values represent grey matter (4-9), which is divided into laminar components, with more myelinated layers located deeper in the cortex (t-distributions 4,5,…,9 are termed T1 layers 6,5,…,1 respectively)

##### 4. Cortical volume sampling and composition analysis:

One of the main challenges in implementing spherical sampling of low-resolution data lies in the resolution difference between the spherical sampling system (average radius of ~0.5 mm) and the T1 data ( $0.67^3\ mm^3$). Overcoming this resolution gap demands a sampling solution based on estimation of subvoxel information (similarly to Shamir et al. 2019). Our solution includes the following steps:

a. Partitioning each voxel into subvoxels:
Each voxel was partitioned into virtual subvoxels encompassing its entire volume. Degree of partitioning was chosen so that each subvoxel has the largest volume while still enabling accurate group representation of a spherical volume with a radius of ~0.5 mm. Consequently, each voxel was partitioned into $10^3$ subvoxels, each assigned location properties, primarily its location inside or outside of a given sphere.

b. Assignment of spherical volume weights:
Each sphere in the sampling system was assigned weights corresponding to each voxel's contribution to its spherical volume. For each sphere, voxel weights were calculated according to the following:

$$W_{voxel_i,sphere_j} = \frac{N_{voxel_i,sphere_j}}{N_{sphere_j}} \quad (3)$$

Where:

$W_{voxel_i,sphere_j}$- Volume weight of voxel i per sphere j.

$N_{voxel_i,sphere_j}$ - Number of subvoxels from voxel i located inside sphere j.

$N_{sphere_j}$ - Total number of subvoxels located inside sphere j.

c. Cortical composition analysis inside sphere:
In order to estimate the cortical composition inside the sampling system, each sphere's volume weights were multiplied by their corresponding voxel's entire probabilistic content (see *T1 probabilistic classification*). In other words, the assignment of spherical weights does not make any assumptions regarding layer order. The process was repeated across all spheres in the entire sampling system, according to the following equation:

$$P(t_k/\text{sphere}) = \sum_{i=1}^{M} \sum_{k=2}^{7} W_{voxel_i,sphere_j} \cdot P(t_k/\text{voxel}_i) \quad (4)$$

Where:
$P(t_k/\text{sphere})$- Probability of t-distribution k per sphere.
k- t-distributions 4,5,6,7, representing T1 layers 4,3,2,1 (respectively).
M- Number of voxels within which sphere j lies.
$W_{voxel_i,sphere_j}$- Volume weight of voxel i per sphere j.
$P(t_k/\text{voxel}_i)$- Probability of t-distribution k in voxel i.

#### 5. DTI analysis and tractography

The DWI dataset was used for global white matter connectivity analysis. Following standard DTI analysis (to extract the FA, MD as well as the fiber orientation density function for each voxel), constrained spherical deconvolution (Tournier 2007) was implemented for estimation of white matter fiber orientation distribution, using ExploreDTI (Leemans et al. 2009). The resulting whole-brain tractography was then analyzed for connectivity across FVE91 regions (see figure 4).

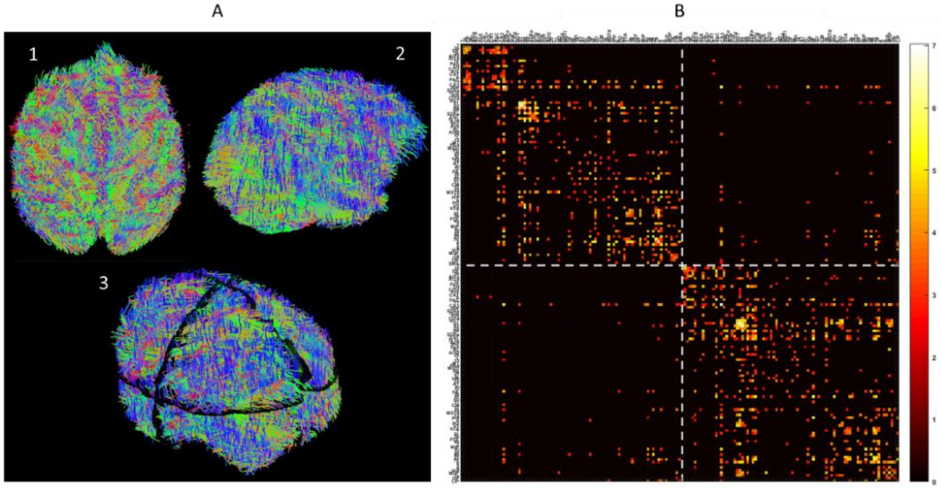


**Fig. 4** Macaque cortical connectivity:

A- Whole brain white matter tractography: top view (1), right side view (2), occipital view (3), visualized using ExploreDTI (Leemans et al. 2009);

B- Connectivity matrix between FVE91 atlas regions, represented as log(number of tracts)

#### Cortical granularity indices

An atlas of cortical granularity indices was formed based on a map of cytoarchitectonic features across the primate cortex (Beul and Hilgetag 2019). The chosen map features the variation in neuron density across M132 atlas regions of the macaque brain. The values, presented as the number of neurons per squared millimeter, were converted to a discrete scale of granularity indices, where higher ranges of neuron densities correlate to higher granularity indices (and vice versa). Missing values were interpolated using an average of the closest existing region values. Indices were then converted to the FVE91 atlas space, using the most common index (see figure 5).

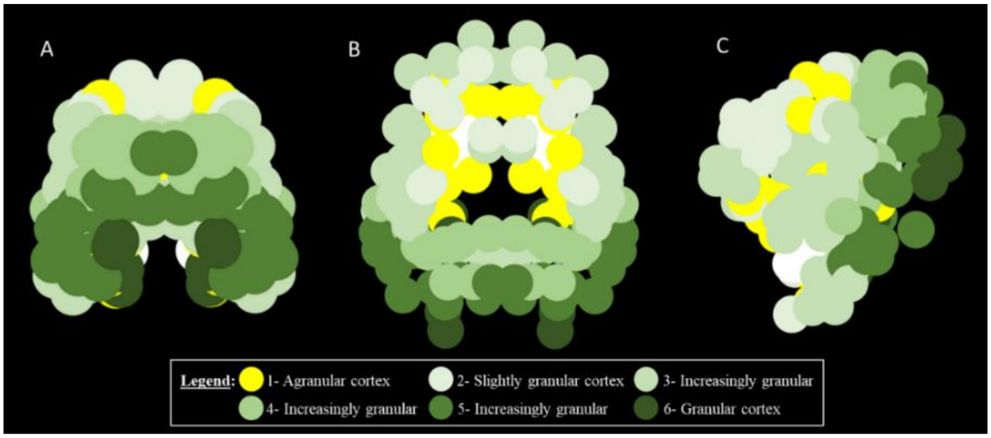


**Fig. 5** Granularity indices across FVE91 atlas regions:
A- Occipital view; B- top view; C- left side view of left hemisphere

### Results

Following are the supplementary figures for the Results and Discussion sections (subsection numbering corresponds to manuscript).

#### Standard cortical connectivity

For an adapted, digitized version of Felleman and Van Essen's reported connections between 35 regions of the visual cortex, see figure 6 (below).

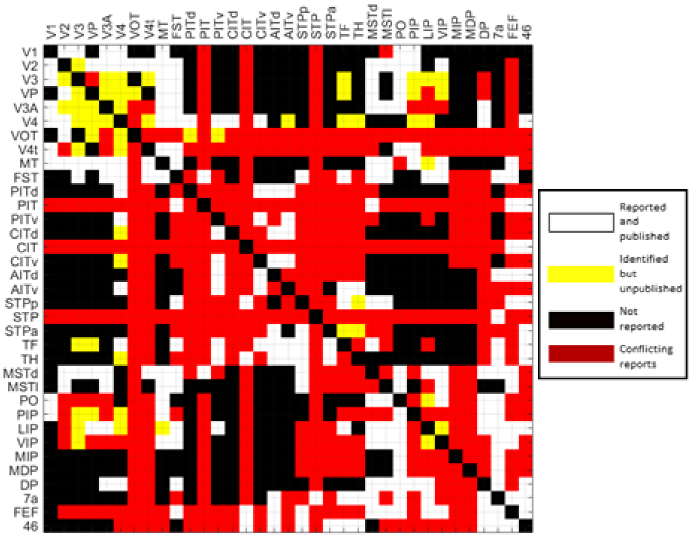


**Fig. 6** Connectivity matrix of cortical connections reported through tracing
(adapted from Felleman and Van Essen 1991)

DTI tractography in the visual cortex was compared to Felleman and Van Essen's reported connections between 35 regions of the visual cortex. For a dendrogram plot of the hierarchical relationship between regions in the macaque visual cortex, see figure 7 (below).

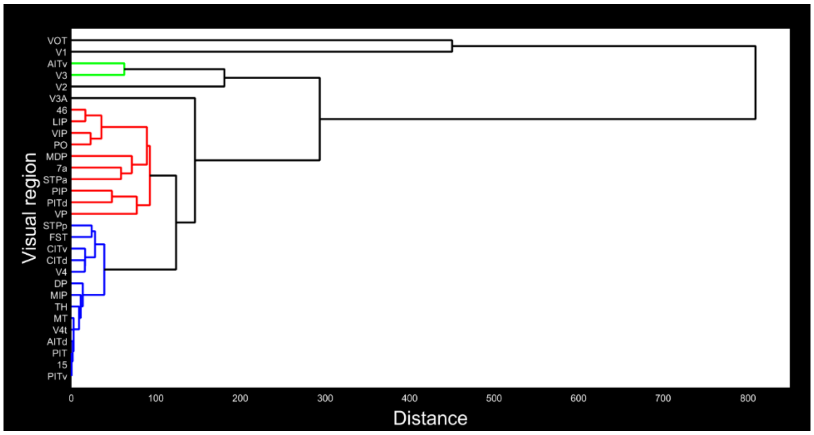


**Fig. 7** Hierarchical relationship between regions in the macaque visual cortex:
A dendrogram plot of the hierarchical cluster tree, representing hierarchical groups of interconnected regions in the macaque visual cortex; Agglomerative clustering was completed using the inner squared distance (minimum variance algorithm) method

In order to quantitatively compare the results in the visual cortex, we binarized both connectivity matrices (see figure 8). We used Felleman and Van Essen's reported connectivity as the actual results, and our tractography connectivity as the predicted results. Consequently, the accuracy value was estimated at just over 71%.

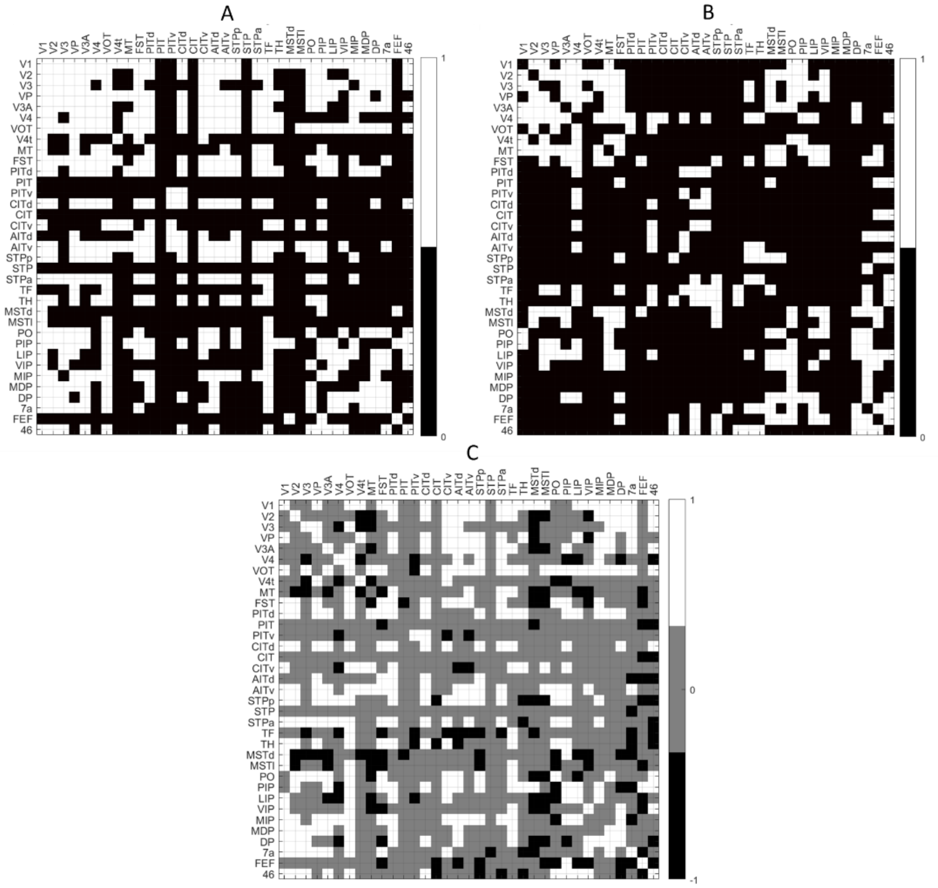


**Fig. 8** Macaque binary connectivity matrices in the visual cortex:

- A- Binary connectivity matrix, representing connections estimated in visual regions through tractography
- B- Binary connectivity matrix, representing cortical connections reported through tracing (adapted from Felleman and Van Essen 1991)
- C- Comparison matrix, representing the difference between matrix A and matrix B

Once the best fiber tracking routine was selected, we used a threshold to remove low connectivity values. The threshold was selected by evaluating positive predictive values (PPVs) for different threshold values across all seven macaque brains (see table 1), followed by averaging of the best performing values and resulting in a threshold of 13.5 tracts.

| Threshold | Positive predictive value (PPV) | | | | | | |
|---|---|---|---|---|---|---|---|
| | (Macaque 1) | Macaque 2 | Macaque 3 | Macaque 4 | Macaque 5 | Macaque 6 | Macaque 7 |
| 0(1) | 0.278 | 0.251 | 0.225 | 0.244 | 0.265 | 0.231 | 0.261 |
| 3 | 0.2806 | 0.255 | 0.239 | 0.250 | 0.270 | 0.243 | 0.247 |
| 5 | **0.283** | **0.261** | 0.238 | **0.259** | 0.258 | 0.246 | 0.251 |
| 10 | 0.282 | 0.257 | 0.228 | 0.259 | 0.266 | 0.231 | **0.262** |
| 20 | 0.250 | 0.251 | **0.252** | 0.243 | 0.286 | **0.256** | 0.252 |
| 30 | 0.259 | 0.246 | 0.275 | 0.254 | **0.293** | 0.248 | 0.235 |

**Table 1** Tractography threshold selection:
Positive predictive values (PPVs) were evaluated for 6 different thresholds: 0 (or 1 tract), 3, 5, 10, 20 and 30, across all seven macaque brains

#### Cortical laminar connectivity

We used the Circular-Connectome toolbox in order to display and explore the resulting model of whole-brain laminar connectivity (best viewed online in 3D). Since the resulting network is multilayered, we expanded each regional node into three laminar positions, represented by a different location in the z axis (lowest circle of nodes- infragranular, mid circle of nodes- granular, and highest circle of nodes- supragranular). In addition, because of the multitude of connections and nodes, we also present intralaminar and interlaminar connections separately (see figure 9).

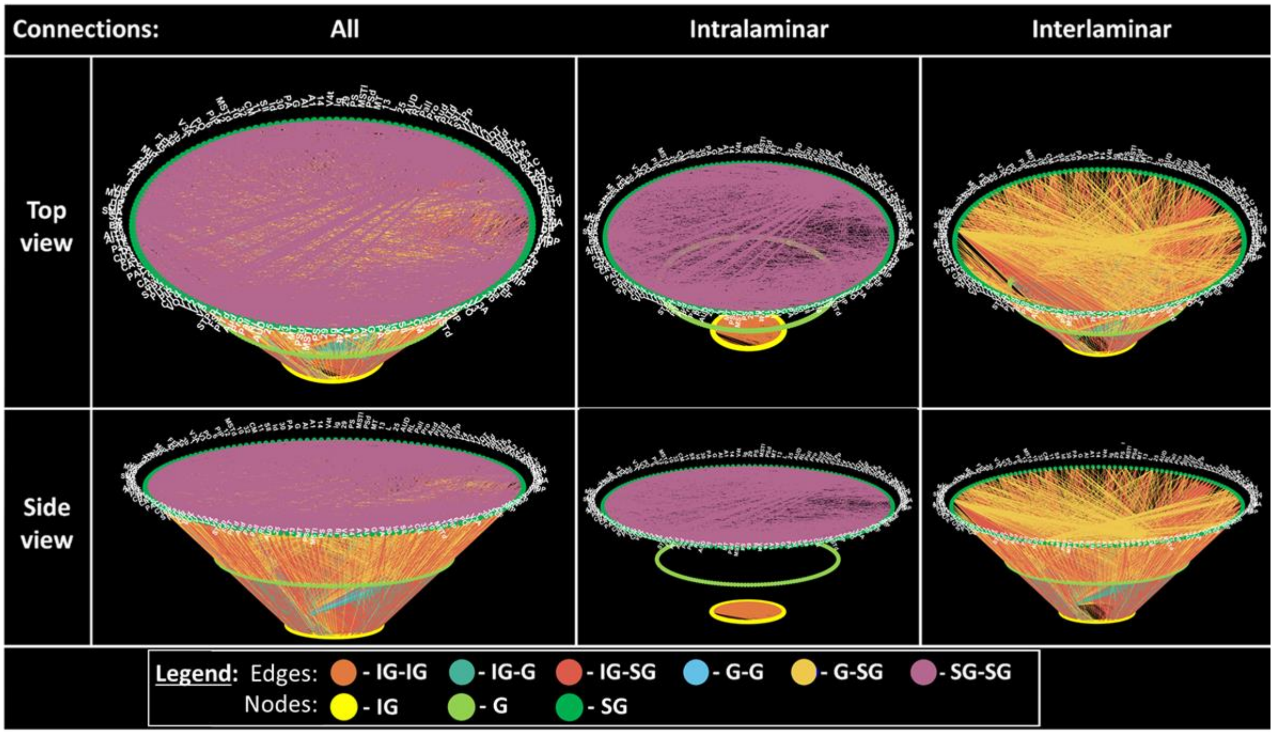


**Fig. 9** A multilayered circular network graph of macaque cortical laminar connectivity:
Columns: first column represents all model connections, second represents connections within a laminar group, and the third represents connections between laminar groups;
Rows: top row includes a top view of the network graph, while the bottom row includes a side view;
Legend: connections were colored according to connection type, based on the connecting laminar groups
(Circular-Connectome toolbox available at: github.com/ittais/Circular-Connectome)

In order to quantitatively compare the results in the visual cortex, once again we binarized both laminar connectivity matrices (see figure 10). Similarly to the standard connectivity comparison, we used Felleman and Van Essen's reported laminar connectivity as the actual results, and our resulting model of cortical laminar connectivity as the predicted results. Consequently, the accuracy value was estimated at just over 83%.

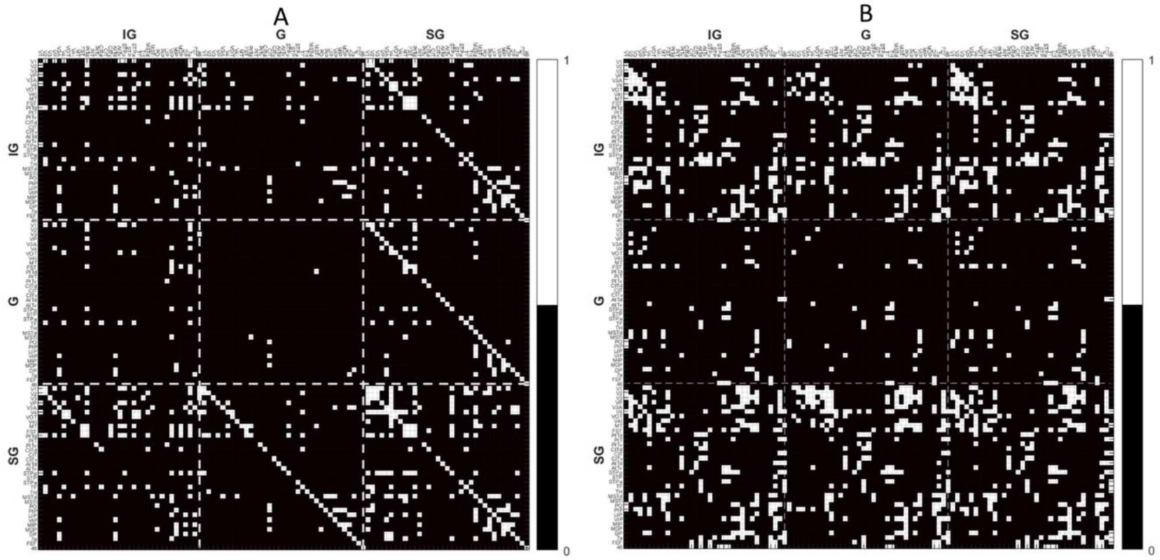


**Fig. 10** Macaque binary laminar connectivity matrices in the visual cortex:

A- Binary supra-adjacency matrix, representing existing laminar-level connections in visual cortex (model results)

B- Binary supra-adjacency matrix, representing cortical laminar connections reported through tracing (a summarized adaptation of connections reported in Felleman and Van Essen 1991)

Where: IG- infragranular, G- granular, SG- supragranular

#### Connectivity results for the 6 additional macaque brains

The connectivity analysis process was repeated for 6 additional excised macaque brains, also from the Mammalian MRI (MaMI) database (Assaf et al. 2020). The T1 layer composition of the original macaque brain was used for all the additional 6 macaque brains. For each macaque brain, connectivity analysis of both standard cortical connectivity and cortical laminar connectivity was conducted. Each resulting connectome was compared in the visual cortex to Felleman and Van Essen's reported findings. Subsequently, the accuracy matrices of both connectomes were compared for each of the six macaques against the exemplary macaque brain (see figure 11 below). The comparison showcases relatively low consistency in a few specific connections, such as MT-LIP, MT-7a and STPp-V3. Nonetheless, overall high consistency can be seen across the macaque brains, specifically in connections between V1, V2 and V3 and across regions such as CIT, AITv and region 46.

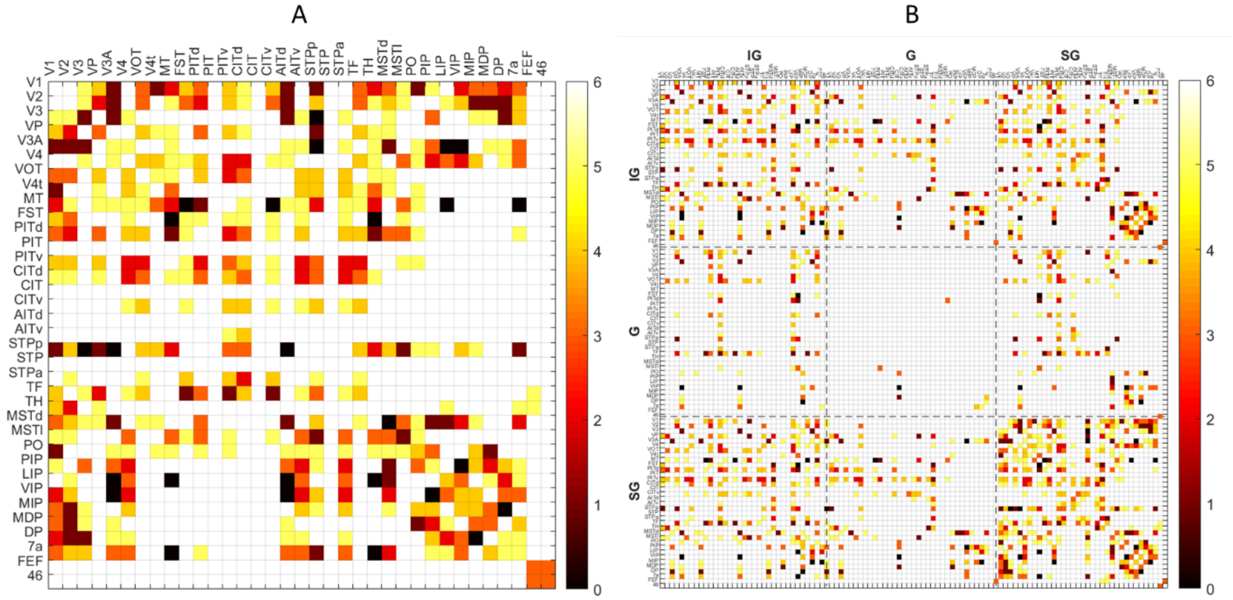


**Fig. 11**: Consistency of connectivity patterns in the visual cortex across macaque brains:
For all macaque brains, accuracy matrices were evaluated of both standard and laminar connectomes in the visual cortex, compared to Felleman and Van Essen's findings. Each of the six macaque brains was then compared to our chosen exemplary macaque brain:

A- Consistency of standard connectivity in the visual cortex
B- Consistency of laminar connectivity in the visual cortex
For both matrices, higher values represent regions (A) or laminar components in regions (B) that are more consistent with the exemplary macaque brain

Patterns of both cortical and cortical laminar connectivity for each macaque were compared to Felleman and Van Essen's 1991 findings in the visual (see table 2).

| # | Macaque | Cortical connectivity (tractography) | | | | | Cortical laminar connectivity (model) | | | | |
|---|---|---|---|---|---|---|---|---|---|---|---|
| | | TPR | TNR | FPR | FNR | Accuracy | TPR | TNR | FPR | FNR | Accuracy |
| (1) | (Macaque1) | 26.03 | 85.74 | 14.26 | 73.97 | 71.51 | 10.8 | 93.53 | 6.47 | 89.2 | 83.32 |
| 2 | Macaque2 | 36.64 | 81.35 | 18.65 | 63.36 | 70.69 | 17.56 | 90.93 | 9.07 | 82.44 | 81.87 |
| 3 | Macaque3 | 21.92 | 86.92 | 13.08 | 78.08 | 71.43 | 13.81 | 93.45 | 6.55 | 86.19 | 83.62 |
| 4 | Macaque4 | 32.19 | 83.17 | 16.83 | 67.81 | 71.02 | 12.2 | 92.98 | 7.02 | 87.8 | 83.01 |
| 5 | Macaque5 | 33.56 | 82.64 | 17.36 | 66.44 | 70.94 | 16.16 | 91.62 | 8.38 | 83.84 | 82.3 |
| 6 | Macaque6 | 21.23 | 87.25 | 12.75 | 78.77 | 71.51 | 9.18 | 94.31 | 5.69 | 90.82 | 83.8 |
| 7 | Macaque7 | 29.45 | 86.6 | 13.4 | 70.55 | 72.98 | 13.45 | 94.01 | 5.99 | 86.55 | 84.06 |
| 7 total | *Average* | 28.72 | 84.81 | 15.19 | 71.28 | 71.44 | 13.31 | 92.98 | 7.02 | 86.69 | 83.14 |
| | *Standard deviation* | 5.46 | 2.2 | 2.2 | 5.46 | 0.69 | 2.7 | 1.16 | 1.16 | 2.7 | 0.74 |

**Table 2** Accuracy evaluation for all 7 macaque brains, including the original macaque brain (1) and the additional 6 brains (2-7):

For each macaque, patterns of both cortical connectivity (left columns) and cortical laminar connectivity (right columns) were compared to Felleman and Van Essen's 1991 findings in the visual cortex; Each comparison includes the following confusion matrix values: true positive rate (TPR), true negative rate (TNR), false positive rate (FPR), false negative rate (FNR), accuracy and F1 score;
Averages and standard deviations of values are shown in the bottom two rows

### Model application on human subject

The entire framework for cortical laminar connectivity was also applied on a human subject. The subject was neurologically and radiologically healthy with no history of neurological diseases. The subject signed informed consent before enrollment in the study. The imaging protocol was approved by the institutional review boards of Sheba Medical Center and Tel Aviv University, where the MRI investigations were performed.

The subject was scanned on a 3T Magnetom Siemens Prisma (Siemens, Erlangen, Germany) scanner with a 64-channel RF coil. One MRI sequence was used to map the cortical connectome (1) and two T1-weighted MRI sequences were used to characterize the cortical layers (2, 3):

1. A standard diffusion-weighted imaging (DWI) sequence, with the following parameters: Δ/δ=60/15.5 ms, b max=5000 (0 250 1000 3000 & 5000) s/mm2, with 87 gradient directions, FoV 204 mm, maxG= 7.2, TR=5200 ms, TE=118 ms, voxel size 1.5×1.5×1.5 $mm^3$, image size 128×128×94 voxels. This sequence was used for global white matter connectivity analysis.
2. An MPRAGE sequence, with the following parameters: TR/TE = 1750/2.6 ms, TI = 900 ms, voxel size 1×1×1 $\boldsymbol{mm^3}$, image size 224×224×160 voxels, each voxel fitted with a single T1 value. This sequence was used as an anatomical reference with high gray/white matter contrast.
3. An inversion recovery echo planar imaging (IR EPI) sequence, with the following parameters: TR/TE = 10,000/30 ms and 60 inversion times spread between 50 ms up to 3,000 ms, voxel size 3×3×3 $\boldsymbol{mm^3}$, image size 68×68×42 voxels, each voxel fitted with up to 7 T1 values (Lifshits et al. 2018). The acquisition time for the inversion recovery data set was approximately 12 min. This sequence was used for cortical laminar composition analysis.

#### Global white matter connectivity analysis

DWI datasets were analyzed for global white matter connectivity using constrained spherical deconvolution (CSD) in ExploreDTI (Leemans et al. 2009). The resulting connectome represents whole-brain cortical connectivity (see figure 12).

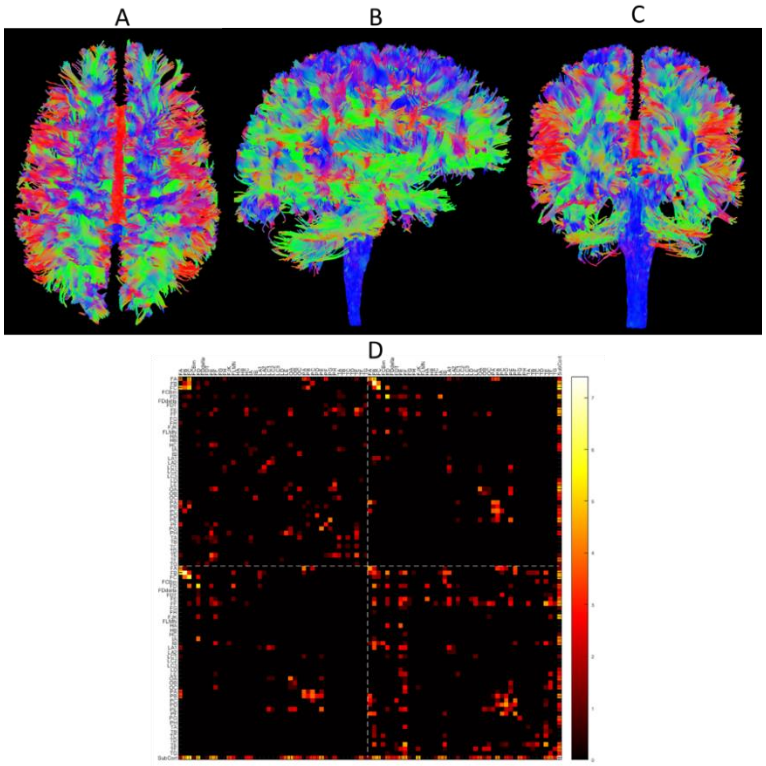


**Fig. 12** Cortical connectivity:

Top- Whole brain white matter tractography: top view (A), right side view (B), occipital view (C), visualized using ExploreDTI (Leemans et al. 2009);
Bottom- Connectivity matrix between Von Economo- Koskinas atlas regions (D), represented as log(number of tracts)

#### Cortical laminar composition analysis

The MPRAGE and IR EPI datasets were analyzed for cortical laminar composition (using the framework presented in Shamir et al. 2019). For the T1 probabilistic classification to tissue types and the resulting cortical laminar probability maps, see figure 13.

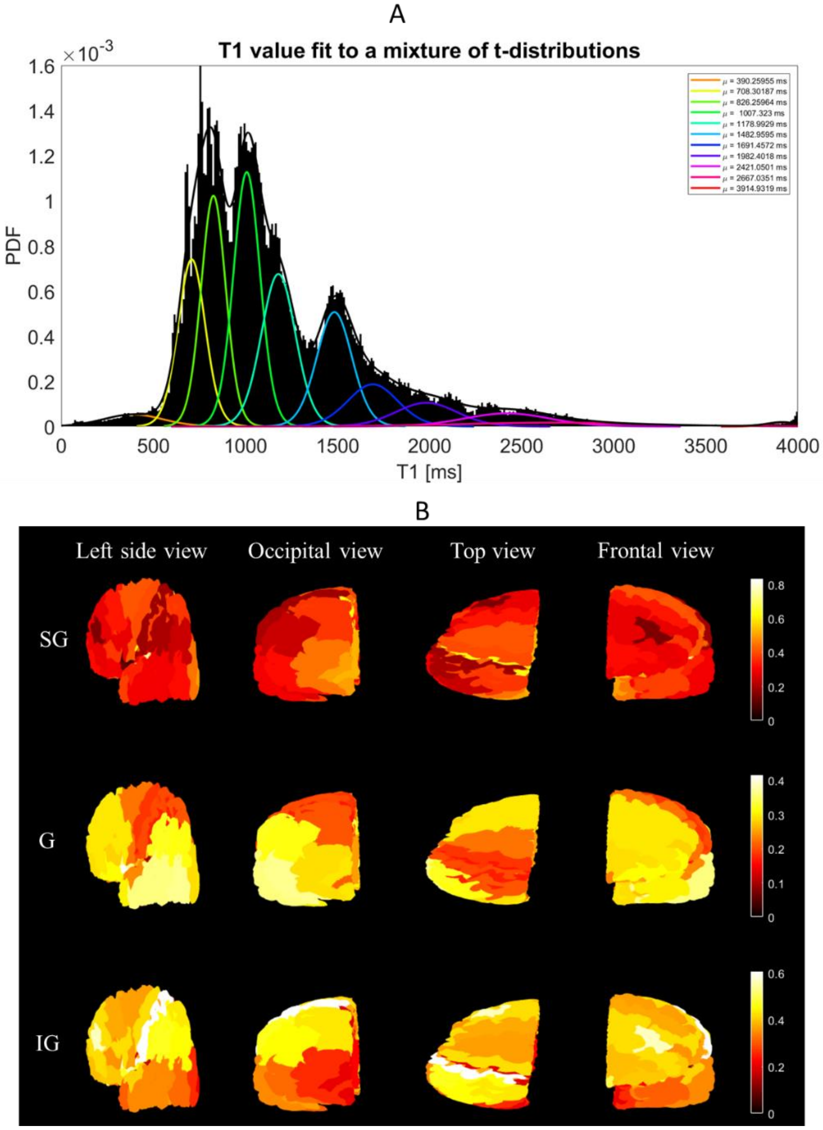


**Fig. 13** Cortical laminar composition analysis:

A- A histogram of T1 values (black) is fit to a mixture of 11 t-distributions (color map). Low T1 values represent white matter (1-3), high values represent fluid/noise (10-11), and mid values represent grey matter (4-9), which is divided into laminar components, with more myelinated layers located deeper in the cortex (t-distributions 4,5,…,9 are termed T1 layers 6,5,…,1 respectively)

B- Cortical laminar probability maps across the left hemisphere (scale according to each range of values):
Top row: supragranular (SG) composition, corresponding to T1 layers 1-3;
Mid row: granular (G) composition, corresponding to T1 layer 4;
Bottom row: infragranular (IG) composition, corresponding to T1 layers 5-6;

#### Model of cortical laminar connectivity

##### Model input:

For a full visualization of all model input datasets in the Von Economo- Koskinas atlas space, including global cortical connectivity, cortical laminar composition and granularity indices, see figure 14 (below).

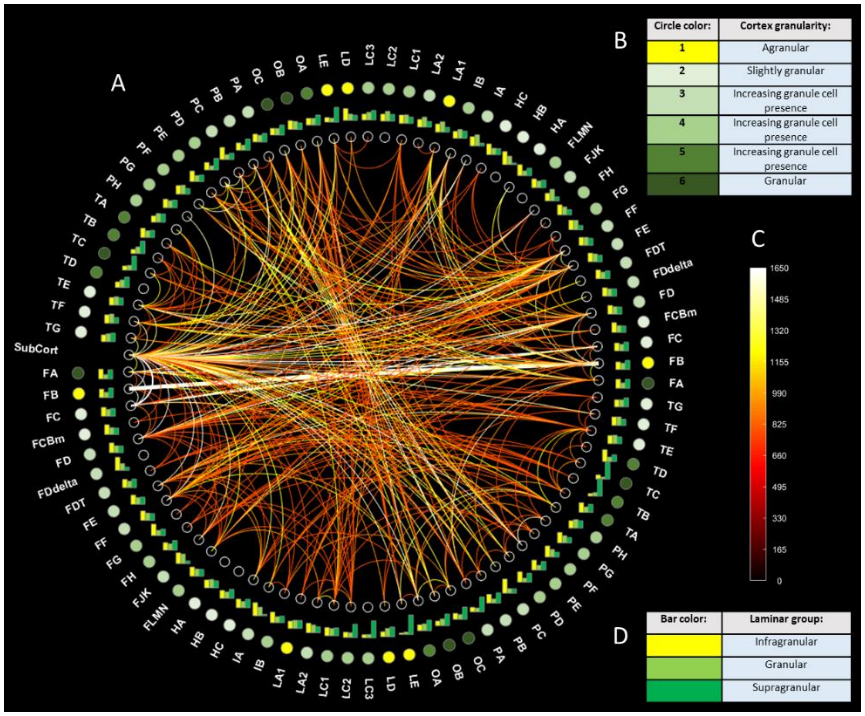


**Fig. 14** Cortical connectivity:

A circular network graph of all model input datasets (A), including:

B- Legend for granularity indices, located in the outermost circle;

C- Color scale for global white matter connectomics, representing connection strength of edges;

D- Legend for color bars, representing grey matter laminar composition, located in the first outer circle;

(visualized using Circular-Connectome toolbox, available at: github.com/ittais/Circular-Connectome)

All input datasets were used in order to apply our model of cortical laminar connectivity (Shamir and Assaf 2020).

**Model output:**

Once again, we used the Circular-Connectome toolbox in order to display and explore the resulting model of whole-brain laminar connectivity. For a full visualization of the resulting model of cortical laminar connectivity, see figure 15 (below).

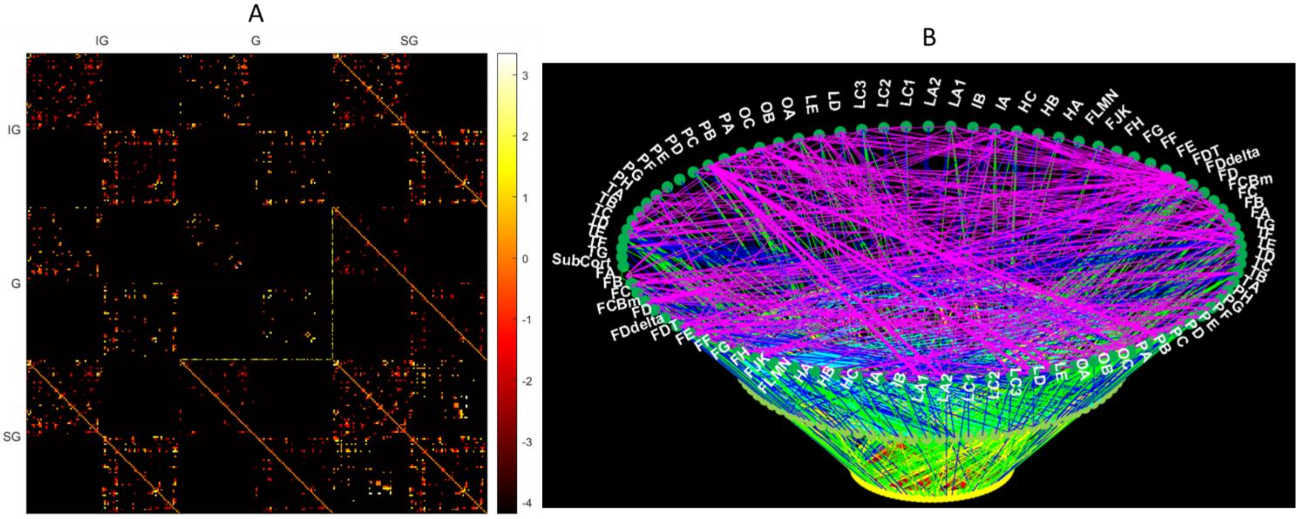


**Fig. 15** Cortical laminar connectivity:

A- Supra-adjacency matrix, representing whole-brain laminar-level connections across Von Economo- Koskinas atlas regions (model results, displayed as log(number of tracts) )

B- A multilayered circular network graph representing cortical laminar connectivity, with connections colored according to connection type, based on the connecting laminar groups
(Circular-Connectome toolbox available at: [github.com/ittais/Circular-Connectome](https://github.com/ittais/Circular-Connectome))

We explored both the standard connectome as well as the laminar connectome using complex tools network analysis tools, similarly to the way the macaque connectomes were explored. The results of the analysis, including the different criteria explored, can be seen in figure 16 (below).

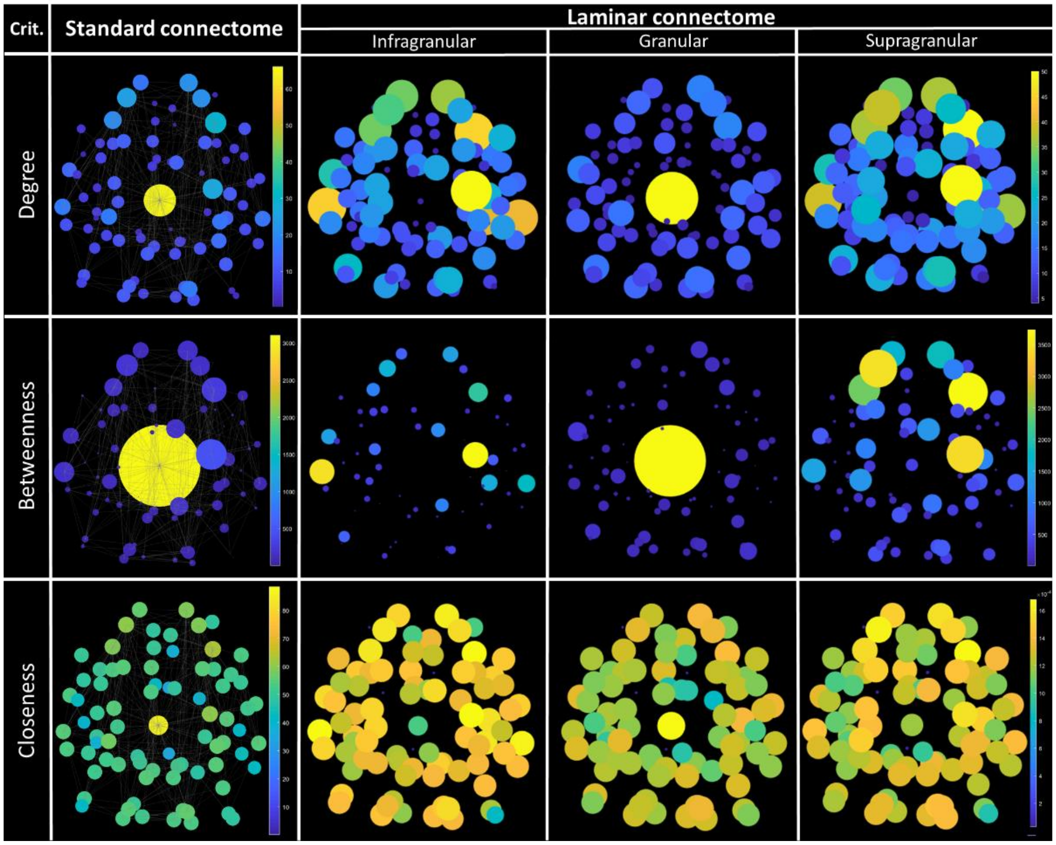


**Fig. 16** Comparison of network criteria of standard versus laminar connectomes:

Columns: first column represents standard connectome; second, third and fourth columns represent laminar connectome components (infragranular, granular and supragranular components respectively);

Rows: top row includes node degrees, mid row includes node betweenness and bottom row includes node closeness (all represented across FVE91 regions);

All images from a top view, with node values scaling both circle size and color (legend on right)
